# Sparse Machine Learning Pipeline with Stabl Identifies Cord Blood Multi-Omic Signatures of Bronchopulmonary Dysplasia

**DOI:** 10.64898/2026.09.12.748996

**Authors:** Karen K. Mestan, Janu Newar, Jiaqi Zhao, Abhik Chakraborty, Jonathan Reiss, William Funk, Ina Stelzer, Benjamin Waked, Gregoire Bellan, Xavier Durand, Julien Hedou

**Author notes:** **Corresponding Author:** Karen K. Mestan, M.S., M.D. 9500 Gilman, M/C 0760, La Jolla, CA 92093.

## Abstract

**Background:** Several omics studies have been completed in recent years, with the goal of identifying biomarkers of complex multifactorial diseases, such as bronchopulmonary dysplasia (BPD).

**Objective:** To evaluate the performance of 3 distinct omics platforms, using a machine learning pipeline with integration of sparse, reliable and adaptive biomarker identification (Stabl).

**Methods:** Using a well-characterized birth cohort, cord blood metabolomics, proteomics and adductomics data were integrated with Least Absolute Shrinkage and Selection Operator (LASSO) regression and Stabl, to evaluate predictive performance for BPD.

**Results:** Sparse multivariable modeling of 45,000 features measured in 217 infants (52 term, 165 extremely preterm ≤28 weeks; 82 with BPD and 35 with severe BPD/death) identified a perfect signature for preterm birth with both LASSO and Stabl (AUROC=1.0; p<0.001). Analysis of the preterm group yielded excellent predictive power for severe BPD (AUROC=0.83; p=0.005). Stabl identified a set of 12 biomarkers (2 adducts, 3 proteins and 7 metabolites) with good performance for predicting grade III BPD (AUROC=0.76; P=0.03). Biomarkers across the 3 omics platforms revealed dysregulated pathways of innate/adaptive immune responses, metabolic programming and oxidative stress.

**Conclusions:** The sparse machine learning pipeline is a complementary approach for identifying novel pathways and biomarkers of multifactorial BPD and its endotypes.

**Impact Statement:**

- Cord blood multi-omics provides a snapshot at birth of the complex interplay between the perinatal metabolome, proteome and exposome.
- We leveraged novel machine learning approaches including integration of sparse, reliable and adaptive biomarker identification (Stabl) to identify novel cord blood biomarkers that elucidate perinatal pathways of BPD, mediated by preterm birth.
- Biomarkers across 3 omics platforms revealed dysregulated pathways of innate/adaptive immune responses, metabolic programming and oxidative stress.
- The sparse machine learning pipeline is a complementary approach for identifying novel biomarkers of multifactorial BPD and its endotypes.

## INTRODUCTION

Bronchopulmonary dysplasia remains the most common chronic lung complication of preterm birth. In recent years, there have been considerable efforts to leverage technologic advances in the -omics to identify the pathogenic pathways and biomarkers that can help predict BPD and also inform the management of premature infants at highest risk. Among these platforms, metabolomics has been the most widely studied, with emerging insights into the role of early metabolic programming in BPD pathogenesis.^1–7^ The development of high-plex mass spectrometry and aptamer-based proteomics has allowed investigation of a wide range of immune and non-immune features in plasma, leading to the discovery of novel proteins and their interactions within a single spot of blood.^8–10^ We have recently applied cord blood adductomics—the study of addition products (adducts)—which serve as exposure biomarkers circulating at birth to investigate the role of intrinsic and extrinsic inflammation, stress and toxicant exposures (i.e., the perinatal exposome) in the development of BPD and other outcomes of preterm birth.^11^

With the growing number of studies leveraging these unprecedently large omics platforms, the next important steps will be to conduct multi-omics analysis in which these features are integrated to further understand the complex interplay and relative importance of distinct -omics datasets. Multi-omics analysis of the outcomes of preterm birth, such as BPD, is limited by patient sample size, variations in timing, volume and availability of specimens collected and the number of features measured. While many of these limitations are being addressed by recent technologic and computational advances, a major barrier remains the cost. These and other limitations have hindered advancement in the field of neonatal-perinatal medicine.

The objective of this study was to evaluate promising and novel multi-omics pipelines that have the potential to overcome the above challenges. Leveraging a well-characterized birth cohort with a large subsample of extremely preterm infants with and without BPD, we utilized high-content omic technologies across 3 available cord blood datasets (metabolomics, proteomics and adductomics), coupled with standard and advanced computational methods, to identify candidate biomarkers of BPD. We employed Stabl, a generalized machine learning method that identifies a sparse, reliable set of biomarkers by integrating noise injection and data-driven signal-to-noise thresholds into multivariable predictive modeling.^12^ With this combined dataset of *over 45,000 features*, we sought to determine the performance of these analytic pipelines in addressing the following 3 questions: 1) Is there a global immune or non-immune signatures associated with preterm birth? 2) Among infants born preterm, can these signatures predict BPD? 3) Can these findings be extended to predict distinct endotypes of BPD linked to BPD severity?

## METHODS

### Study Design and Patient Cohort

A total of 217 infants were included in the study, with multi-omics profiling performed across three data layers: metabolomics, proteomics and adductomics. The patient sample was drawn from a single-center cohort study conducted in Chicago (Prentice Women’s Hospital), with ongoing enrollment since 2008. The details of the parent cohort have been previously published.^11,13–17^

**Figure 1A-1C** shows the distribution of patients and number of features in each of the 3 datasets. The total number of infants and their corresponding cord blood samples included in the analysis was 217, taking account substantial overlap as well as patients who were included in at least one of the 3 omics datasets. The adductomics dataset included 210 patients and 105 features (comprising 56 known and 49 unknown compounds). The details of this dataset and the adductomics features identified have been previously published.^11^ The metabolomics dataset consisted of 209 patients with 39,038 features identified, while the proteomics dataset included 50 patients with 6,432 quantified proteins. Cord blood plasma untargeted metabolomics was completed by Sapient BioAnalytics, LLC (San Diego, CA) using rLC-MS. Aptamer-based proteomics was completed by Somalogics (Boulder, CO) on 50 cord blood samples using the SomaScan 7K Assay. Each of the above 3 platforms were completed in single runs: Adductomics in 2022, metabolomics in 2023 and proteomics in 2024.

**Figure 1:**
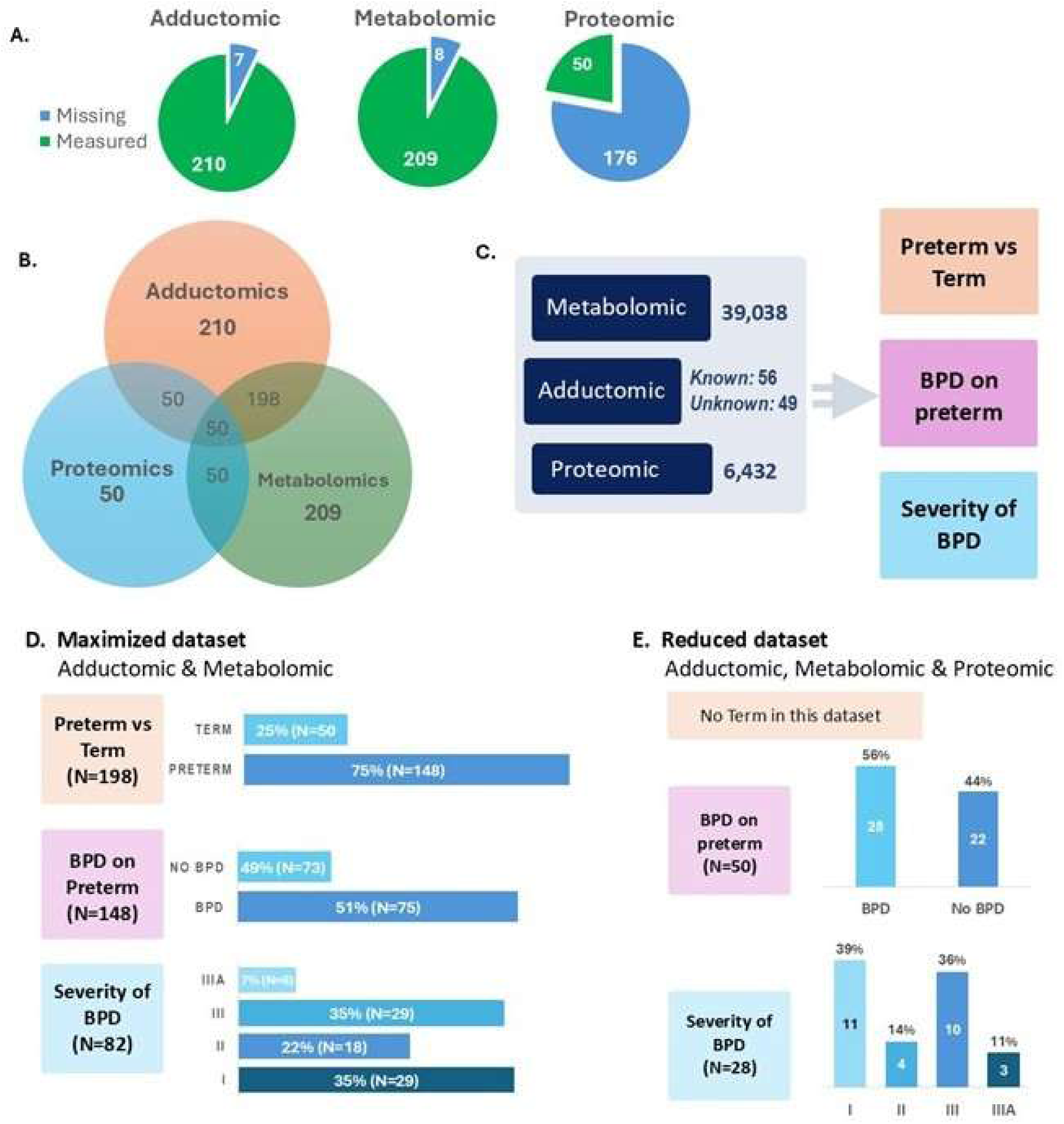
Project overview showing distribution of the total study sample, which includes 217 patients with cord blood biomarker data from any one or all three of the omics datasets: (A) Pie charts demonstrating the proportion of the total patient sample with completed (green) and not completed (blue) omics platforms. Due to cost, proteomics was only completed on a subset of extremely preterm infants. (B) Venn diagram of the number of patients and overlap with data available for each omic type; (C) Overview of datasets, number of features and main outcomes; (D) Distribution of patients in the “maximized dataset” in which only adductomics and metabolomics were completed; (E) Distribution of patients in the “reduced dataset” in which only a subset of patients were profiled, but all 50 patients had metabolomics, adductomics and proteomics completed.

### Clinical Outcomes Assessment

The total study sample consisted of full-term births and extremely preterm births, which allowed us to determine, at baseline, how cord blood multi-omic signatures vary between the two extremes of the gestational age spectrum. Gestational age at delivery, confirmed by last menstrual period (LMP) and early ultrasound before 20 weeks gestation, was used to classify infants as term (born ≥37 weeks) or preterm (≤28 completed weeks gestation). This binary outcome was assessed across 2 of the 3 omics datasets (adductomics, metabolomics) to identify molecular and immune signatures associated with extremely preterm delivery.

Among the preterm births (≤28 weeks completed weeks), BPD was studied as the primary outcome. Infants were categorized as either BPD-positive (Grade I, II, III or IIIA) or BPD-negative (non-BPD), based on the 2018 NICHD Workshop Criteria.^18^ A total of 165 preterm infants were included in this analysis to explore predictors of disease occurrence. Within the subset of infants diagnosed with BPD (N=82), disease severity was further stratified according to Grade I, II, III or IIIA BPD. This outcome was analyzed to determine whether molecular and immune variables could discriminate between different severity levels of lung disease.

Two analytic strategies were used to include the entire sample of 217 patients, taking into account completeness of the omics datasets, but also overlap and inclusion of all 3 omics. To achieve this, a “maximized dataset” was analyzed, which included all study subjects with both adductomics and metabolomics completed (N=198) (**Figure 1D**). In order to include all 3 omics, we also analyzed a “reduced dataset” which included all study subjects in which all 3 omics were completed (N=50) (**Figure 1E**). Since a major limiting factor in the proteomics project was cost, only a subset of the preterm infants had proteomics completed. That study was designed to directly compare BPD versus gestational-age and sex-matched non-BPD infants. Thus, the reduced dataset was well-matched and restricted to a narrower gestational age range.

### Data Preprocessing and Multi-omics integration

Each omics dataset was preprocessed individually prior to integration. For the proteomic data, a log₂ transformation was applied to stabilize variance and approximate normality across protein abundance values. A standardized preprocessing pipeline was then applied to each dataset independently. This ensured consistent data quality across omics layers: 1) *<u>Variance Threshold</u>* removed features with zero variance (uninformative variables); 2) *<u>Low Info Filter</u>* excluded features with excessive missingness (>20% missing values); 3) *<u>Simple Imputer</u>* replaced remaining missing values using the median of each feature; 4) *<u>Standard Scaler</u>* standardized all features using Z-score normalization to ensure comparability across scales.

For multi-omics integration, two complementary strategies were used. The first strategy was to maximize the number of patients in the dataset by including those with at least 2 complete omics datasets. N=198 patients met these criteria in which both adductomics and the metabolomics were completed (**Figure 1D**). The second strategy was to use all the omics possible by including all patients with all 3 omics datasets completed. This led to a reduced the number of patients to 50 in which metabolomics, adductomics and proteomics were completed (**Figure 1E**).

### Missing data handling

To minimize the need for data imputation, analyses were performed using participants with complete data (following the two approaches mentioned before) for the relevant omics layers and outcomes. Each dataset underwent individual preprocessing, during which features with more than 20% missing values were removed using the LowInfoFilter step. Remaining missing values were imputed using the median within each dataset.

### Statistical Analysis

A comprehensive list of maternal and infant covariables known to be associated with preterm birth and BPD were followed and recorded in the larger parent cohort. The list of relevant covariates analyzed is shown in **Table 1**. Univariate analyses were performed to assess the association between each covariate and the clinical outcomes. For binary outcomes, the Mann–Whitney U-test was applied to compare distributions between groups. To adjust for multiple comparisons and control the false discovery rate, the Benjamini–Hochberg procedure was employed.^19^

**Table 1.** Baseline demographics and clinical characteristics of births included in the analysis.

|  | **Maximized Dataset**<br>**N=217** |  | **P** | **Reduced Dataset**<br>**N=50** |  | **P** |
| --- | --- | --- | --- | --- | --- | --- |
|  | **Full-Term**<br>**N=52** | **Preterm**<br>**N=165** |  | **No BPD**<br>**N=22** | **BPD**<br>**N=28** |  |
| **Gestational age (weeks)<sup>a</sup>**<br>**mean± SD** | 39.5 ± 0.1 | 27.0± 0.1 | <0.001 | 27.6 ± 0.2 | 26.5 ± 0.3 | 0.003 |
| **Birth weight (grams)** | 3427.7± 62.3 | 991.0 ± 21.3 | <0.001 | 1088.0± 30.1 | 915.5 ± 34.8 | 0.001 |
| **Birth weight percentile** | 52.8 ± 3.6 | 60.2± 2.0 | 0.073 | 67.3 ± 3.4 | 60.4 ± 4.9 | 0.281 |
| **Maternal age (years)** | 34.7± 0.6 | 31.0 ± 0.4 | <0.001 | 30.8 ± 1.1 | 32.3 ± 1.1 | 0.322 |
| **Apgar 1-min,**<br>**median [IQR]** | 8 [8, 9] | 5 [3, 7] | <0.001 | 6 [6, 7] | 4 [3, 6] | 0.001 |
| **Apgar 5-min** | 9 [9, 9] | 7 [6, 8] | <0.001 | 8 [6, 8] | 6 [5, 8] | 0.016 |
| **Preeclampsia, n (%)** |  |  |  |  |  |  |
| no | 51 (98) | 139 (84) | 0.007 | 18 (82) | 23 (82) | 0.976 |
| yes | 1 (2) | 26 (16) |  | 4 (18) | 5 (18) |  |
| **Chorioamnionitis** |  |  |  |  |  |  |
| no | 50 (96) | 146 (88) | 0.175 | 20 (91) | 24 (86) | 0.683 |
| yes | 2 (4) | 19 (12) |  | 2 (9) | 4 (14) |  |
| **Infant sex** |  |  |  |  |  |  |
| male | 26 (50) | 83 (50) | 0.970 | 9 (41) | 16 (57) | 0.254 |
| female | 26 (50) | 82 (50) |  | 13 (59) | 12 (43) |  |
| **Gestation Type:** |  |  |  |  |  |  |
| singleton | 50 (96) | 117 (71) | <0.001 | 12 (55) | 21 (75) | 0.171 |
| twin | 2 (4) | 46 (28) |  | 9 (41) | 7 (25) |  |
| triplet | 0 (0) | 2 (1) |  | 1 (4) | 0 (0) |  |
| **Membrane Rupture:** |  |  |  |  |  |  |
| SROM | 10 (19) | 86 (52) | <0.001 | 10 (45) | 13 (46) | 0.945 |
| AROM | 42 (81) | 79 (48) |  | 12 (55) | 15 (54) |  |
| **Mode of Delivery:** |  |  |  |  |  |  |
| Vaginal | 45(87) | 62 (38) | <0.001 | 10 (45) | 9 (32) | 0.336 |
| C-section | 7 (13) | 103 (62) |  | 12 (55) | 19 (68) |  |
| **GBS Status:** |  |  |  |  |  |  |
| neg | 28 (54) | 65 (39) | <0.001 | 15 (68) | 9 (32) | 0.025 |
| pos | 19 (36) | 24 (15) |  | 3 (14) | 4 (14) |  |
| unknown | 5 (10) | 76 (46) |  | 4 (18) | 15 (54) |  |
| **Antenatal Steroids:** |  |  |  |  |  |  |
| complete | 0 (0) | 130 (79) | <0.001 | 22 (100) | 17 (61) | 0.001 |
| incomplete | 0 (0) | 27 (16) |  | 0 (0) | 9 (32) |  |
| none | 52 (100) | 8 (5) |  | 0 (0) | 2 (7) |  |
| **Preterm Labor:** |  |  |  |  |  |  |
| no | 52 (100) | 50 (30) | <0.001 | 4 (18) | 6 (21) | 0.776 |
| yes | 0 (0) | 115 (70) |  | 18 (82) | 22 (79) |  |
| **Prolonged ROM:** |  |  |  |  |  |  |
| no | 52 (100) | 84 (51) | <0.001 | 12 (55) | 17 (61) | 0.661 |
| yes | 0 (0) | 81 (49) |  | 10 (45) | 11 (39) |  |
| **Preterm Premature ROM:** |  |  |  |  |  |  |
| no | 49 (94) | 113 (68) | <0.001 | 17 (77) | 23 (82) | 0.669 |
| yes | 3 (6) | 52 (32) |  | 5 (23) | 5 (18) |  |
| **Abruption**<br>**no**<br>**yes** | 52 (100)<br>0 (0) | 145<br>20 | 0.005 | 20 (91)<br>2 (9) | 28 (100)<br>0 (0) | 0.189 |
| **Maternal Race:** |  |  |  |  |  |  |
| **White** | 40 (77) | 61 (37) | <0.001 | 8 (36) | 11 (39) | 0.944 |
| **Black/African American** | 3 (6) | 42 (25) |  | 6 (27) | 7 (25) |  |
| **Asian** | 2 (4) | 13 (8) |  | 2 (9) | 3 (11) |  |
| **Other** | 5 (9) | 32 (19) |  | 5 (23) | 4 (14) |  |
| **Unknown** | 2 (4) | 17 (10) |  | 1 (5) | 3 (11) |  |
| **Ethnicity:** |  |  |  |  |  |  |
| **Not Hispanic or Latino** | 42 (81) | 115 (70) | 0.043 | 15 (68) | 23 (82) | 0.511 |
| **Hispanic or Latino** | 10 (19) | 35 (21) |  | 5 (23) | 4 (14) |  |
| **Unknown** | 0 (0) | 15 (9) |  | 2 (9) | 1 (4) |  |
| **BPD Outcomes:** |  |  |  |  |  |  |
| **Any BPD<sup>b</sup>** |  |  |  |  |  |  |
| **No** | 217 (52) | 83 (50) | <0.001 | 22 | 0 | <0.001 |
| **Yes** | 0 (0) | 82 (50) |  | 0 | 28 |  |
| **None** | 52 (100) | 83 (50) | <0.001 | 22 (100) | 0 (0) | <0.001 |
| **Grade I** | 0 (0) | 29 (18) |  | 0 (0) | 11 (39) |  |
| **Grade II** | 0 (0) | 18 (11) |  | 0 (0) | 4 (14) |  |
| **Grade III** | 0 (0) | 29 (17) |  | 0 (0) | 10 (36) |  |
| **Grade IIIA** | 0 (0) | 6 (4) |  | 0 (0) | 3 (11) |  |

<sup>a</sup> Gestational age at birth confirmed by last menstrual period (LMP) and early ultrasound before 20 weeks gestation. <sup>b</sup> BPD (yes) defined as having Grade I, II, III, or IIIA as determined at 36 weeks postmenstrual age by NICHD Workshop criteria (Higgins, et al. 2018).

### Multivariate Modeling

A 5×5 Monte Carlo cross-validation (MCCV) scheme was implemented for model evaluation. The dataset was randomly partitioned into five folds, repeated five times to reduce partition-induced bias. Predictions for each patient were aggregated by taking the median value across all folds.

The performance metric for the continuous outcome was the root mean squared error (RMSE) to account for the error magnitude. For binary outcomes, the area under the receiver operating characteristic curve (AUROC) was calculated to assess the model’s ability to distinguish between positive and negative classes. Statistical significance of the models was assessed using the Mann–Whitney U-test for classification tasks and Pearson’s correlation coefficient for regression tasks, with a significance level set at p<0.05.

### Modeling Strategy using advanced machine learning process

Our primary predictive model was Lasso^20^ regression with L1 regularization, used to promote sparsity and perform feature selection during cross-validation. In the context of multi-omics data integration, we implemented a late fusion strategy. Separate Lasso models were trained on each individual proteomics panel dataset within the cross-validation framework, and their predictions were subsequently aggregated using a stacked generalization approach. This allowed us to leverage the unique predictive power of each omic layer while combining them into a cohesive predictive model. The performance of these models was evaluated based on cross-validated metrics to assess their predictive capability.

### Biomarker Identification with Stabl

To identify stable and sparse sets of biomarkers, we used the Stabl machine learning framework as first described by Hedou, et al.^12^ Stabl was applied after cross-validation and trained on the entire dataset to extract relevant features. Stabl operates without prior assumptions by introducing artificial features into the original dataset, creating an augmented dataset. This dataset undergoes regularization methods like Lasso over multiple bootstrap iterations, resulting in a feature selection frequency graph. A reliability threshold is established to distinguish informative real features from artificial ones, optimized to minimize the inclusion of artificial features while maximizing the retention of relevant biomarkers.

By analyzing the frequency with which each variable was selected across the bootstrap iterations, we assessed the robustness and importance of each feature. This approach enabled us to identify variables with consistent predictive value, providing insights into potential biological mechanisms underlying BPD.

For multi-omics data integration with Stabl, a halfway fusion strategy was adopted. Feature selection was performed individually on each omic dataset using Stabl, and the selected features were then combined into a single dataset for final analysis. This method facilitated the identification of predictive signatures associated with clinical outcomes.

### Confounder Analysis

To account for potential confounders, we integrated relevant covariates (**Table 1**) into multivariable regression models. The primary model examined the relationship between biological predictors and clinical outcomes, while the extended model combined the primary model’s predictions with the identified confounders to assess their influence on the outcomes. We analyzed changes in the estimated coefficients and tested for statistical significance to evaluate the impact of confounding, considering it significant when changes in model predictions showed a statistically meaningful p-value. This analysis aimed to determine how the associations observed were independent of these potential confounding factors, thereby strengthening the validity of the findings.

## RESULTS

### Study Participant Characteristics

A total of 217 cord blood samples were included in this study. Multi-omics profiling was performed on clinical data encompassing three omics layers: adductomics, metabolomics, and proteomics. The adductomics dataset was comprised of 210 samples with 105 detected features (56 known and 49 unknown). The metabolomic dataset included 209 samples with 39,038 features, while the proteomic dataset included 50 cord blood samples with 6,432 features. The distribution of outcomes (preterm versus full-term; No BPD versus BPD) is displayed in **Table 1** for both the maximized and reduced datasets.

### Multi-omics Analysis

**Table 2** summarizes the AUROC for each combined and individual omics analysis for the maximized and reduced datasets. **Figure 2** illustrates the predictive performance of the LASSO and Stabl models through Receiver Operating Characteristics (ROC) for each of the 6 analyses, which are summarized below:

**Figure 2:**
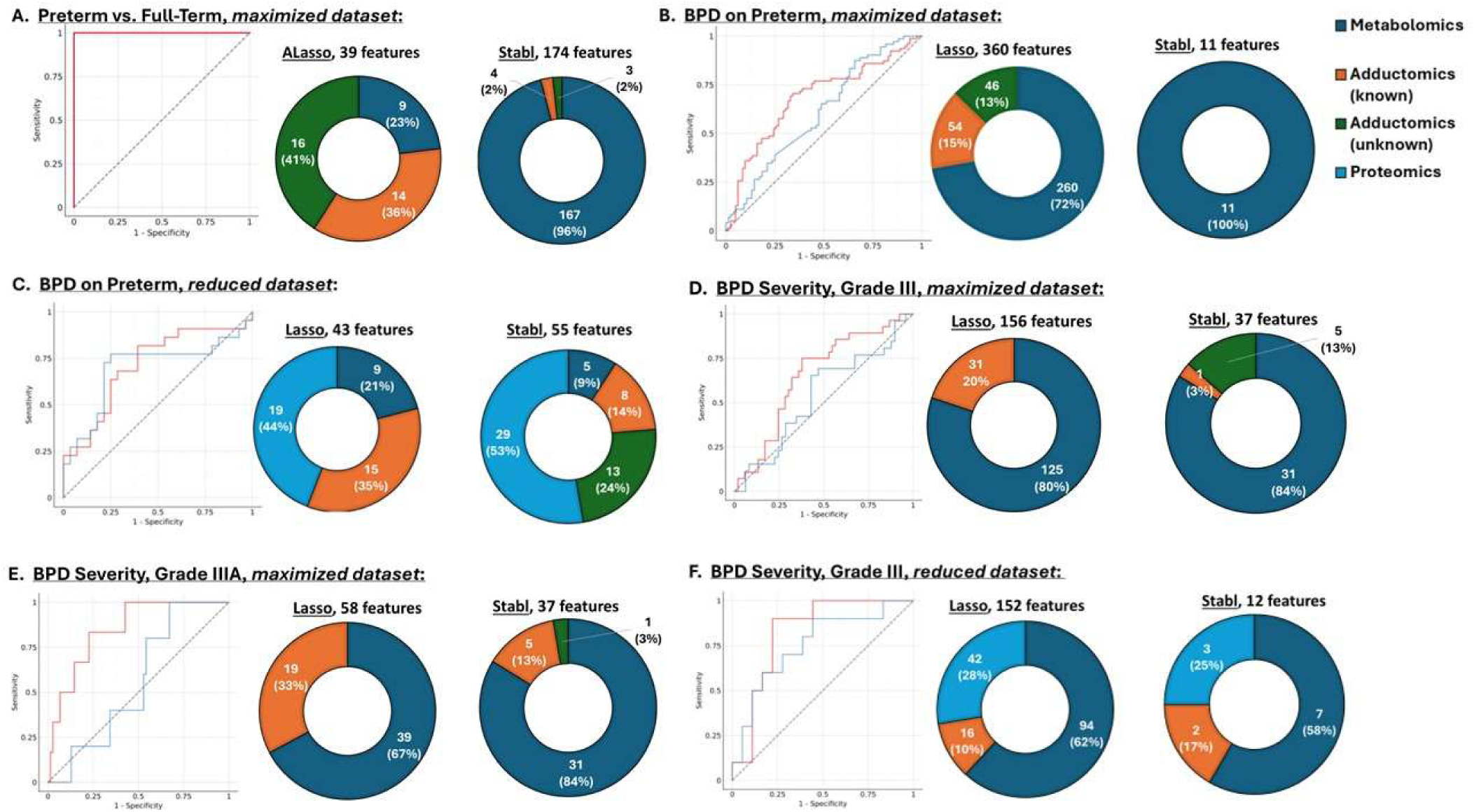
Model performances and distribution of selected features of the multi-omics analyses. **(A) Analysis for Preterm Birth.** The performance of the ALASSO model is illustrated through a Receiver Operating Characteristic (ROC) curve (thick red line), achieving an Area Under the Curve (AUC) of 1.00, indicating a perfect predictive performance (thick red line). The straight dashed line represents the observation of a random model with no predictive power. Pie charts illustrate the distribution of features by ALasso (left) and further by Stabl (right). Dark blue—metabolomic features; Orange—known adductomics features; Green—unknown adductomics features. **(B) Analysis for BPD on preterm in the maximized dataset.** Performances of the LASSO model (red line) and Stabl (blue line) are illustrated through ROC curves. The straight dashed line represents the observation of a random model with no predictive power. Pie charts illustrate the distribution of features by LASSO (left) and Stabl (right). Stable selected fewer features overall that were all metabolites (dark blue). **(C) Analysis of BPD (yes/no) with the reduced dataset.** The performance of the models are illustrated through ROC curves for LASSO (red line) and Stabl (blue line). The straight dashed line represents the observation of a random model with no predictive power. Pie charts illustrate the % distribution of selected features, with prominence of proteomic features (light blue) with the reduced dataset. **(D) Analysis for BPD Severity (Grade III) with the maximized dataset.** The performance of the LASSO model is illustrated through ROC curves for LASSO (red line) and Stabl (blue line). The straight dashed line represents the observation of a random model with no predictive power. Pie charts demonstrate the predominance of metabolomic features (dark blue) selected by both LASSO and Stabl, and more representation of unknown adducts (green) by Stabl. **(E) Analysis for BPD Severity (Grade IIIA) with the maximized dataset.** Relative model performances and distribution of features are similar to Grade III of the maximized dataset. **(F) Analysis for BPD Grade III with the reduced dataset.** Performance of the models are illustrated through ROC curves for LASSO (red line) and Stabl (blue line). Pie charts illustrate the % distribution of selected features, with prominence of proteomic features (light blue) with the reduced dataset. Stabl narrowed the number of selected features to 12 biomarkers, represented by metabolomics (58%), adductomics (17%) and proteomics (25%).

**Table 2.** Summary of Model Performance and Number of Selected Omic Features.

| Model | Omics | AUROC (95% CI) | # of Selected Features |
| --- | --- | --- | --- |
| ***Analysis for Preterm vs. Full-term Birth (maximized dataset)*** |  |  |  |
| ALasso | Metabolomics | 1.00 (95% CI: 1.00, 1.00) *** | 9 |
| ALasso | Adductomics (known) | 0.85 (95% CI: 0.79, 0.91) *** | 14 |
| ALasso | Adductomics (unknown) | 0.87 (95% CI: 0.80, 0.92) *** | 16 |
| ALasso | All | 1.00 (95% CI: 1.00, 1.00) *** | 39 |
| **Stabl** | **All** | **1.00 (95% CI: 1.00, 1.00) \*\*\*** | **174** |
| ***Analysis for BPD on Preterm (maximized dataset)*** |  |  |  |
| LASSO | Metabolomics | 0.66 (95% CI: 0.57, 0.75) *** | 260 |
| LASSO | Adductomics (known) | 0.50 (95% CI: 0.41, 0.59) * | 54 |
| LASSO | Adductomics (unknown) | 0.65 (95% CI: 0.57, 0.74) *** | 48 |
| LASSO | All | 0.68 (95% CI: 0.59, 0.76) *** | 360 |
| **Stabl** | **All** | **0.61 (95% CI: 0.52, 0.69) \*** | **11** |
| ***Analysis of BPD on Preterm (reduced dataset)*** |  |  |  |
| ALasso | Metabolomics | 0.67 (95% CI: 0.52, 0.81) * | 9 |
| ALasso | Adductomics (known) | 0.63 (95% CI: 0.46, 0.80) | 15 |
| ALasso | Adductomics (unknown) | 0.50 (95% CI: 0.50, 0.50) | 0 |
| ALasso | Proteomics | 0.55 (95% CI: 0.38, 0.71) | 19 |
| ALasso | All | 0.71 (95% CI: 0.56, 0.86) ** | 43 |
| **Stabl** | **All** | **0.70 (95% CI: 0.53, 0.85) \*\*** | **55** |
| ***Analysis of BPD Severity (maximized dataset)*** |  |  |  |
| ***Grade III BPD versus rest:*** |  |  |  |
| LASSO | Metabolomics | 0.66 (95% CI: 0.53, 0.78) * | 125 |
| LASSO | Adductomics (known) | 0.58 (95% CI: 0.44, 0.72) | 31 |
| LASSO | Adductomics (unknown) | 0.50 (95% CI: 0.50, 0.50) | 0 |
| LASSO | All | 0.66 (95% CI: 0.53, 0.78) * | 156 |
| **Stabl** | **All** | **0.54 (95% CI: 0.41, 0.68)** | **37** |
| ***Grade IIIA BPD versus rest:*** |  |  |  |
| LASSO | Metabolomics | 0.77 (95% CI: 0.48, 0.98) * | 39 |
| LASSO | Adductomics (known) | 0.60 (95% CI: 0.19, 0.93) | 19 |
| LASSO | Adductomics (unknown) | 0.50 (95% CI: 0.50, 0.50) | 0 |
| LASSO | All | 0.85 (95% CI: 0.70, 0.97) ** | 58 |
| **Stabl** | **All** | **0.56 (95% CI: 0.35, 0.80)** | **37** |
| ***Analysis of BPD Severity, Grade III versus rest (reduced dataset)*** |  |  |  |
| LASSO | Metabolomics | 0.63 (95% CI: 0.41, 0.82) | 94 |
| LASSO | Adductomics (known) | 0.73 (95% CI: 0.52, 0.91) | 16 |
| LASSO | Adductomics (unknown) | 0.50 (95% CI: 0.50, 0.50) | 0 |
| LASSO | Proteomics | 0.82 (95% CI: 0.64, 0.96) ** | 42 |
| LASSO | All | 0.83 (95% CI: 0.66, 0.96) ** | 152 |
| **Stabl** | **All** | **0.76 (95% CI: 0.56, 0.94) \*** | **12** |

\*P<0.05; \*\*P<0.01; \*\*\*P<0.001

#### Sparse multivariable model identifies a perfect signature predictive of preterm birth

To first gain understanding of the influence of gestational immaturity at birth, multi-omics analysis comparing the extremely preterm to full-term births was performed using the maximized dataset, which included adductomics and metabolomic profiles from the majority of patients (N=198) across both omics layers. Patients were classified according to gestational age at delivery as either preterm (in this dataset, all preterm were ≤28 completed weeks gestation) or term (≥37 weeks), resulting in an unbalanced distribution, with 148 preterm and 50 term infants included in the analysis. To capture potential interactions between features, multivariate machine learning models were trained in a cross-validation framework (see Methods). The adaptive LASSO (ALasso) model, which incorporated data from adductomics and metabolomics, demonstrated perfect predictive performance with an AUROC of 1.00 (95% CI: 1.00, 1.00, p = 1e-16) (**Figure 2A**) with 39 selected features. Models trained on a single panel suggest signal stemming from all the omics with a predominance for metabolomics, with outstanding predictive power (AUROC = 1.0, 95% CI: 1.0–1.0, p<0.001) and selection of 9 biomarkers.

Adductomics (Known) showed excellent predictive power (AUROC = 0.85), with selection of 14 biomarkers. Adductomics (Unknown) also demonstrated excellent predictive power (AUROC = 0.87) with selection of 16 biomarkers. The output of features selected by ALasso and Stabl, with overlap, is provided in **Data Supplement File A**. The majority of metabolomic features remain un-annotated at the time of this analysis. Mass-to-charge ratio (m/z), retention time (RT) and related parameters for each metabolomic and adductomics features are listed in the raw data files through 10.6084/m9.figshare.33147644.

Confounding analysis (see Methods) was performed on the ALasso model (AUROC=1.00) to control for different clinical features while predicting preterm birth. As illustrated in **Data Supplement File B**, several covariates contribute to explain preterm birth, however predictive power of the model remained significant (p<0.001). As expected, clinical features such as birthweight, gestational age and preterm labor showed significant (P<0.05) associations with the outcomes, indicating that part of the model’s predictive signal are driven by underlying clinical characteristics rather than purely molecular signatures.

#### Fair to moderate multivariate predictive power for BPD on preterm with the maximized dataset

Given that full-term and extremely preterm cord blood signatures were expected to be different, the outcome of BPD was analyzed on the maximized dataset of preterm infants only. This resulted in a well-balanced distribution, in which 75 BPD infants and 73 non-BPD infants had both metabolomics and adductomics completed. Next, we trained predictive models to evaluate how well different sets of biomarkers could distinguish between individuals with BPD. The best-performing model was the LASSO model, which incorporated data from both omics. This model achieved a fair predictive performance, with an AUROC of 0.68 (p=7e-5) (**Figure 2B**). Models based on individual omics performed similarly in the fair/moderate range, all P<0.05) (**Table 2**).

To achieve the highest level of prediction, the LASSO model selected 360 biomarkers spanning all omics: 260 from metabolomics, 54 from the known adductomics, and 48 from the unknown adductomics. To further refine biomarker selection, we applied Stabl, our stability-driven selection method which offers a more robust and rigorous feature selection approach. Stabl identified a smaller set of 11 biomarkers (**Figure 2B**). Despite inclusion of both metabolomics and adductomics (known and unknown), Stabl’s stringent selection process resulted in only features from metabolomics being retained. Those biomarkers achieved a fair predictive performance of AUROC = 0.61 (p = 2e-2). A list of LASSO and Stabl selected markers, with overlap is included in **Data Supplement File A**.

Confounding analysis was performed on the LASSO model, using AUROC=0.68 to control for different clinical variables while predicting BPD outcome with the maximized dataset. As illustrated in **Data Supplement File B**, four covariates significantly (p<0.05) contribute to BPD while predictions of the model become non-significant (p=0.24). These included birthweight, unknown maternal race, multiple gestation and rupture of membrane status (artificial versus spontaneous).

#### Good predictive power of BPD on Preterm with the reduced dataset

Since multi-omic analysis of the maximized dataset was limited in that only 2 omics datasets were considered, we applied a second strategy in which all 3 omics (metabolomics, adductomics and proteomics) were completed. Since only a subset of the extremely preterm infant subgroup had proteomics completed, this substantially reduced the sample size. However, this subsample was also better matched by gestational age, birth weight and infant sex, due the original study design of the single proteomics project. This “reduced dataset” was comprised of 50 samples, in which 28 infants were diagnosed with BPD and 22 infants did not develop the condition (non-BPD).

The best predictive model was obtained using ALasso which selected 43 features to explain BPD in preterm infants. Statistical analysis revealed good predictive power for those features with an AUROC of 0.71 (p = 1e-2) (**Figure 2C**). The predictive power based on single omics (**Table 2**) ranged from fair/moderate for metabolomics (AUROC=0.67; P<0.05) to non-significant with adductomics and proteomics. Further analysis with Stabl revealed similar predictive power as LASSO (AUROC=0.70; 95% CI: 0.53, 0.85, p=1e-2), but with more selected features and enhanced representation of proteomics (53%) and adductomics (unknown, 24%) (see **Figure 2C**).

Confounding analysis was performed on the LASSO model using AUROC=0.71 to control for different confounders while predicting BPD with the reduced dataset. As illustrated in **Data Supplement File B**, five covariates (gestational age, rupture of membrane type, 1-minute Apgar score, antenatal steroids and birthweight) contribute to explain the outcome at the P<0.05 threshold, while predictions of the model become non-significant (p=0.23).

#### Fair to excellent predictive power for BPD Severity with the maximized dataset

We know that analysis by the dichotomous outcome of BPD (yes/no) does not adequately capture disease heterogeneity—mostly commonly assessed by differences in BPD severity—that could potentially be predicted by the different omics pathways. Thus, we next evaluated whether comparison of the omics according to BPD severity might result in enhanced performance of the multi-omic models. Among the 82 BPD patients, 6 patients for Grade IIIA, 29 for Grade III, 18 for Grade II and 29 for Grade I BPD. To analyze these subgroups, we transformed the multi-grade outcome as several binary outcomes to predict one grade against the other. This resulted in no predictive power for Grade I BPD vs rest and Grade II BPD vs rest in the maximized dataset.

For Grade III BPD vs rest, a fair predictive power was obtained using the LASSO model with an AUROC=0.66 (p = 2e-2) (**Figure 2D**). 156 features were selected by LASSO, while Stabl selected 37 features representing metabolomics, as well as known and unknown adductomics. Similar results were obtained with single omics on metabolomics (AUROC = 0.66, p < 0.05) with 125 selected features.

Analysis of Grade IIIA BPD vs rest showed excellent predictive power with 58 selected features using the LASSO model (AUROC=0.85; p = 3e-3) (**Figure 2E**). Variable predictive power was obtained in single omics analysis (**Table 2**), with metabolomics demonstrating good predictive power (AUROC = 0.77; p < 0.05) with 39 selected features, and adductomics (Known) having fair predictive power (AUROC = 0.60; p > 0.05) with 19 selected features.

In contrast to patterns with the reduced dataset analysis involving proteomics, multi-omic analysis with Stabl in these maximized datasets yielded no significant predictive power, with 37 selected features for both Grade III and IIIA. It is interesting to note that LASSO identified almost identical features for metabolomics in the Grade III vs rest and Grade IIIA vs rest analyses, but the metabolomic features further selected by Stabl were different between III and IIIA BPD (**Data Supplement File A**). Unknown adducts identified by both LASSO and Stabl were identical but with different fold-changes.

Confounding analysis was performed on the LASSO model using AUROC=0.66 for Grade III BPD vs rest and AUROC=0.85 for Grade IIIA BPD vs rest to control for different clinical variables while predicting BPD severity. As illustrated in **Data Supplement File B**, some covariates significantly (p<0.05) contribute to explain the outcome while predictions of the model are not significant (p=0.11) anymore for Grade III BPD vs rest, but remained significant (p=0.01) for Grade IIIA BPD vs rest.

#### Excellent predictive power for BPD Severity with the reduced dataset

Finally, we investigated how analysis of the reduced dataset, taking into account all 3 omics, would perform in identifying selected features predictive of BPD severity. In this analysis, there were only 28 BPD patients with all 3 omics: 11 patients with Grade I, 4 for Grade II, 10 for Grade III and 3 with Grade IIIA BPD. Despite the reduced sample size, we achieved excellent predictive power with the LASSO model for the Grade III (AUROC=0.83; p=5e-3) with 152 selected features (**Figure 2F**). Single metabolomics and adductomics (Known) had moderate and good predictive power, respectively, while single proteomics had excellent predictive power (AUROC=0.82; p<0.01) with 42 selected features (**Table 2**).

To refine biomarker selection further, we applied the Stabl algorithm, which identified a smaller set of 12 key biomarkers while still maintaining good predictive performance (AUROC=0.76; p=3e-2). This set of biomarkers included 7 features from metabolomics (58%), 2 features from adductomics (17%) and 3 features from proteomics (25%). **Table 3** describes the 12 biomarkers selected by Stabl. **Figure 3** illustrates the stability path graphs generated for the distinct omics platforms.

**Figure 3:**
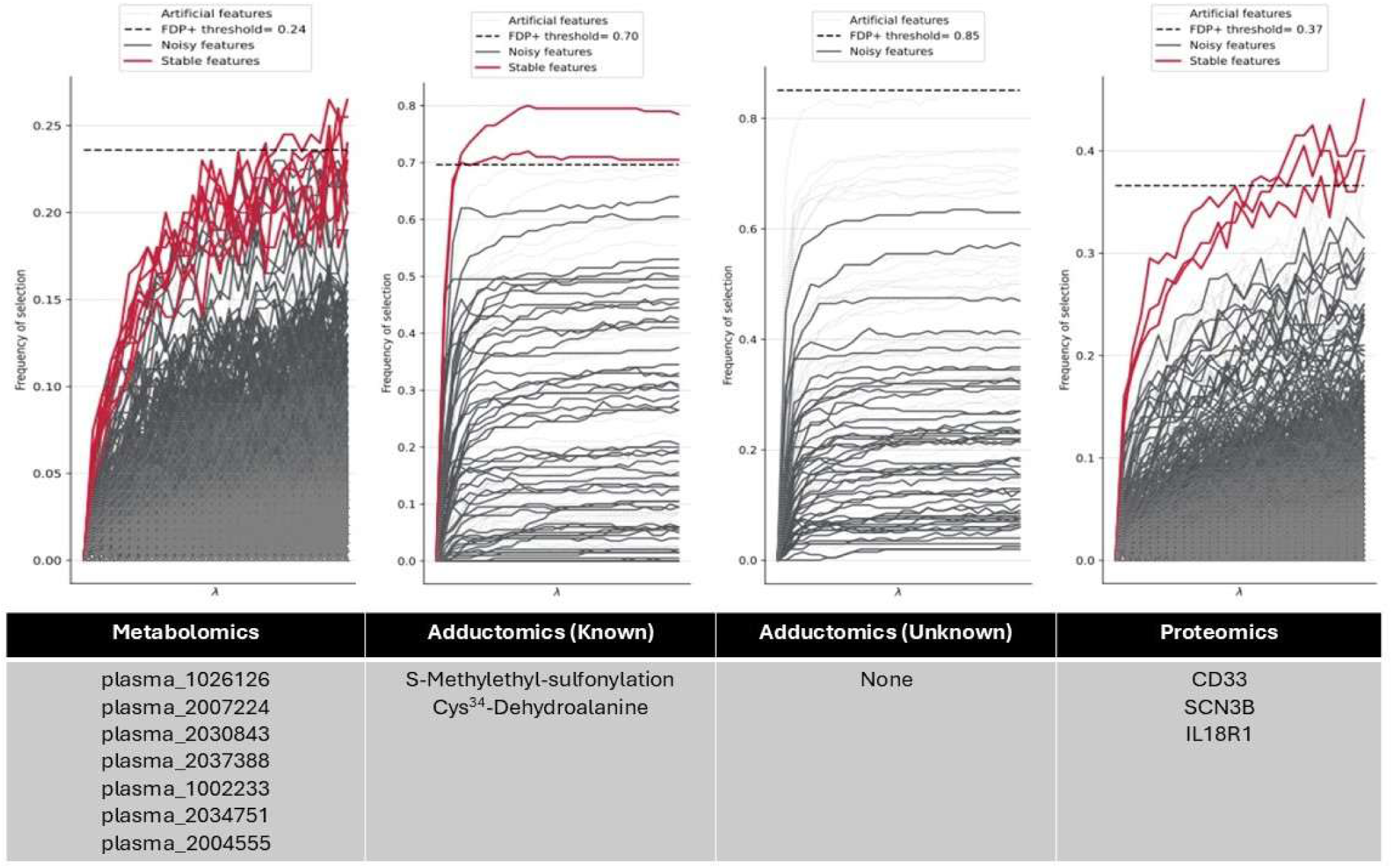
Stability path graphs generated by Stabl for the features selected as predctors of Grade III BPD in the reduced dataset. Each plot shows the relationship between the regularization parameter and selection frequency across bootstrapped samples. Each line represents a variable, with a dashed horizontal line marking a selection frequency threshold for each omics platform. 12 features (listed below each graph) exceeded the respective thresholds for metabolomics, adductomics (known), adductomics (unknown) and proteomics, and can be considered relevant for predicting the patients who develop BPD at 36 weeks postmenstrual age.

**Table 3.** The 12 features selected by Stabl for BPD Grade III with the reduced dataset.

| **Stabl Selected Features** | **Corresponding Omics** | **Annotation** | **Frequency of selection in Cross-Validation** | **Regression Coefficients** |
| --- | --- | --- | --- | --- |
| **A037** | Adductomics | S-Methylethly-sufonylation | 0.93333 | -3.22169 |
| **A003** | Adductomics | Cys<sup>34</sup>→ Dehydroalanine | 0.86667 | -0.15039 |
| **Q9NY72** | Proteomics | SCN3B | 0.53333 | -0.05745 |
| **P20138** | Proteomics | CD33 | 0.46667 | 3.86693 |
| **Q13478** | Proteomics | IL18R1 | 0.33333 | 0.76038 |
| **plasma_1026126** | Metabolomics | Unknown<sup>a</sup> | 0.06667 | 4.71321 |
| **plasma_2004555** | Metabolomics | Unknown | 0.06667 | -1.92992 |
| **plasma_2007224** | Metabolomics | Unknown | 0.06667 | 2.90778 |
| **plasma_2030843** | Metabolomics | Unknown<sup>b</sup> | 0.06667 | 4.69324 |
| **plasma_2034751** | Metabolomics | Unknown<sup>c</sup> | 0.06667 | 2.53686 |
| **plasma_1002233** | Metabolomics | Unknown | 0.0 | 7.12363 |
| **plasma_2037388** | Metabolomics | Unknown<sup>d</sup> | 0.0 | 4.67788 |
| Based upon recent search using the Human Metabolome Database (HMDB) (https://hmdb.ca/spectra/ms/search) by polarity, m/z and RT values (see: [10.6084/m9.figshare.33147644](https://figshare.com/figures/10.6084/m9.figshare.33147644)), unknown features have MSI Level 2 – Level 4 evidence for the following possible metabolites: <sup>a</sup> Feature is either a ceramide or diacylglycerol metabolite; <sup>b</sup> N,N-Diisopropyl-N'-isoamyl-N'-diethylaminoethylurea or Solacaproine; <sup>c</sup> Phosphatidylinositol phosphate (PIP), Phosphatidyl glycerolphosphate (PGP) or Cytidine diphosphate diacylglycerol (CDP-DG); <sup>d</sup> Val-val-tyr-pro-trp-thr-gln-arg-phe (VV-hemorphin-7). |  |  |  |  |

Confounding analysis was performed on the LASSO model using AUROC=0.83 to control for different clinical features while predicting BPD severity. As illustrated in **Data Supplement File B**, five covariates contributed to explain the outcome of Grade III BPD (preterm labor, preeclampsia, maternal age, ethnicity, 1-minute Apgar score) while predictions of the model remained highly significant (p=2.6e-6). Of note, the effects of birth weight and gestational age were non-contributory.

## DISCUSSION

We performed a novel integrated multi-omics analysis of 3 large omics datasets, with over 45,000 features measured in cord blood plasma from a single cohort of preterm and full-term births. We employed a robust pipeline for preprocessing and data integration, taking into account missing data and confounding of a wide range of prenatal, intrapartum and postnatal risk factors. This is the first multi-omics report of its kind to focus on cord blood profiles—which are representative of the perinatal (surrounding birth) metabolome, exposome and proteome across the gestational age spectrum and preterm birth leading up to BPD.

Advanced machine learning and multi-omics integration identified a perfect signature predictive of extremely preterm birth. This signature appeared to be driven by metabolomics, with combined and independent contributions from adductomics—a relatively newer platform that measures relative intensity of addition products (adducts) formed by oxidant stress at birth.^11^ Further analysis of a maximized dataset that included only the preterm subgroup revealed fair multivariate predictive power for BPD, and very good predictive power for BPD severity when all 3 omics datasets were included, despite restriction to a much smaller patient sample (N=50). Specifically, in the reduced dataset in which all 3 -omics data were included, LASSO achieved excellent predictive power for Grade III BPD (i.e., survival with severe disease at 36 weeks), with single proteomics being a top-performer (AUROC=0.82, 95% CI: 0.64-0.96; P<0.01). Application of the Stabl algorithm identified selected features across all omics domains, with ultimately 12 key biomarkers (**Table 3**) identified that maintained good predictive performance for severe BPD.

Several novel patterns of the different models and their relative performances were found, as summarized in **Table 2**. Overall, LASSO with and without Stabl performed similarly across all analyses, in both maximized and reduced datasets, with the exception of Grade IIIA BPD in the maximized datasets. Single omics using LASSO or ALasso revealed variations in predictive power and feature selection among the different omics platforms. With the exception of the largest analysis involving preterm vs. full-term births, Stabl retained or reduced the number of selected features to a smaller but more diverse set of biomarkers by adding subsampling, noise injection and data-driven threshold optimization to improve feature reliability.^12^ This also resulted in better representation by all omics datasets rather than being influenced by the largest group of biomarkers (i.e., metabolomics with 39,000 features). Thus, Stabl may be a preferred approach in analyzing datasets with diverse sets of omics features, even when the patient sample size is relatively small.

The robust confounding analyses in the pipeline revealed multiple covariates that potentially contribute to model performance. This was as expected especially in the maximized cohort that included both full-term and extremely preterm infants. A key finding in the series of these analyses was that after adjustment for these covariates, only 2 approaches preserved significance at the P<0.001 threshold: the preterm vs. full-term birth analysis of the maximized dataset, and grade III BPD vs. rest of the reduced dataset. It is particularly interesting to observe the differences in significant clinical variables among the different analyses, as they may reflect distinct underlying pathophysiological mechanisms and progression stages within the disease spectrum. Such variations suggest that distinct clinical factors drive or modulate biological responses depending on the severity or specific subtype of the condition, offering valuable insights into disease heterogeneity, BPD endotypes and potential stratification strategies.

Another important observation was how the predictive power changed when studying the maximized versus the reduced datasets. Addition of the proteomics dataset, even though the patient sample size was markedly reduced, resulted in preserved predictive performance with Stabl as compared with LASSO only, with the ability to narrow the number of features while taking into account a more diverse biomarker platform that included all 3 omics. Another interesting finding was that the BPD (yes/no) analysis using the maximized dataset, which did not include proteomics or take into account BPD severity, overall did not perform as well with or without Stabl. This was in contrast to the excellent performance of the reduced dataset on BPD severity that included proteomics. Collectively, this suggests that proteomics is an essential key driver of BPD, and particularly for severe BPD. Another important speculation is that the reduction in biomarker “noise” with Stabl, even in the setting of a much smaller sample size, can better identify features of biological importance. This has implications for downstream studies of biomarker characterization upon selection that arise from multi-omics integration.

This is the first report of its kind with a large sample size of term, preterm and BPD infants, a unique focus on cord blood across 3 omics platforms, and an unprecedented number of features and novel combination of omics platforms analyzed. Another favorable aspect is the well-characterized cohort from which the study sample was derived. This allowed us to identify potential covariates of BPD pathogenesis that contribute to the multi-omic profiles. For example, the larger sample of preterm and term births constituted a diverse set of patients in which gestational age, birth weight, mode of delivery, antenatal steroids, maternal race, age and pregnancy complications such as preeclampsia and chorioamnionitis contributed significantly to the “perfect signature” of preterm birth described in our results. In subgroup and reduced datasets, this confounding was minimized considerably down to 5 covariates in the analysis for BPD severity, demonstrating that the analytic pipeline is able to minimize confounding while preserving good predictive potential of the models.

Application of standard and newer analytics to the >45,000 features in cord blood plasma from 217 births resulted in robust comparison of the multivariate models, and selection of promising candidate biomarkers for predicting and understanding preterm birth and BPD. The “perfect signature” achieved in the first analysis confirms that the cord blood samples collected and the omics platforms generated from these samples were of high-quality, to effectively distinguish omics profiles according to the extremes of gestational age at birth. We found that metabolomics and adductomics are key drivers/predictors of extremely preterm birth, while proteomics is a promising platform for understanding and predicting BPD severity. While many of the selected features have yet to be completely annotated, the study demonstrates how the Stabl algorithm can accelerate our investigations by focusing on the most reliably predictive “unknown” biomarkers for future annotation and characterization. For the majority of models, multi-omics approaches outperformed single omics. This has important implications for how we approach future investigations and the development of biomarker panels of meaningful use in the clinical setting.

Metabolomics and proteomics are rapidly growing platforms that have accelerated our research on preterm birth and neonatal outcome. We adopted our approach from Stelzer and colleagues, who used a similar machine learning pipeline to study coordinated alterations in the maternal metabolome, proteome and immunome associated with spontaneous labor, laying the groundwork for developing blood-based methods for predicting day of delivery in preterm and term pregnancies.^21^ Mithal and colleagues used untargeted mass spectrometry to characterize the cord blood proteome across gestational age.^15^ Newer, high-plex aptamer-based proteomics is an emerging platform that allows more comprehensive profiling of up to 11,000 immune and non-immune proteins using very small volumes of plasma. Our study validates that the metabolome and proteome are key predictors of preterm birth outcomes, and is the first reported to integrate untargeted metabolomics and high-plex proteomics for prediction of BPD.

A relatively understudied platform is adductomics, which enables interrogation of the “perinatal exposome” as measured by reactive addition products formed by maternal environmental exposures. These adducts bind to human serum albumin and thus serve as stable biomarkers of peripartum oxidant stress.^22^ In our recently published report on cord blood adductomics,^11^ we identified several adducts associated with preterm birth and BPD. In contrast to that single adductomics analysis, multi-omics integration with metabolomics and proteomics yielded additional selection of adductomic features (see below). Given the higher number and more diverse set of features involved in multi-omics versus individual omics, different profiles and selected features are expected. This highlights the power of multi-omics in predicting and understanding BPD and its endotypes.

The 12 features selected by Stabl (**Table 3**) provide insight into the “perfect storm” of integrated pathways associated with the development and severity of multifactorial BPD. The 7 metabolomic features remain largely unannotated to date. Based upon m/z and RT characteristics, there is MSI/Tier level 2-4 evidence that these selected metabolites are involved in lipid metabolism, specifically lipid compositional changes in the maturing postnatal lung (PIP, PGP, CPD-DG)^23,24^ and regulation of cardiovascular and metabolic disease via inhibition of the renin-angiotensin system (VV-hemorphin-7).^25^ The 3 proteomics features are novel findings: Feature P20138 (positive correlation) was identified by UniProtKB as CD33 (siglec-3), or sialic acid binding immunoglobulin-like lectin 3, which plays a role in cell-cell interaction and immune regulation of monocytes, macrophages and granulocytes, maintaining immune cells in a resting state and suppressing monocyte activation.^26^ Feature Q13478 (positive correlation) was identified as IL-18 Receptor 1 (IL-18R1) which is involved in the adaptive immune and inflammatory responses, specifically involving pathogens associated with intrauterine inflammation.^27,28^ IL18R1 has been reported in association with BPD in African-American infants.^29^ Feature Q9NY72 (negative correlation) was identified by UniProtKB as sodium channel regulatory subunit beta-3 (SCN3B). The relevance of this protein to BPD has not been previously reported or acknowledged.

The 2 adducts identified by Stabl have been annotated: S-methylethyl-sulfonylation and Cys34→Dehydroalanine. Dehydroalanine is a post-translational modification formed when cysteine undergoes oxidative electrophilic attack. In human serum albumin, this modification can occur at Cys34 and affect plasma clearance and enhance free-radical scavenging during oxidative stress.^30^ Relative decreases in S-methylethyl-sulfonylation adducts suggests reduced stable oxidation of cysteine residues, which can be a sign of lower oxidant stress, or altered redox regulation.^31^ These adductomics changes may indicate protective mechanisms or adaptive responses to oxidative challenges.

Limitations of this study are that it only includes 2 and 3 omics platforms, measurement at one timepoint (birth) and in patients from a single birth center. With consideration of each subgroup analysis, the small patient sample size becomes another important limitation. Thus, the generalizability of our findings needs to be validated in future studies involving a more diverse patient population from multiple centers, and analysis of data from serial timepoints and expanded omics platforms. These opportunities for future studies are highly feasible, given the growing number of clinical studies with large omics datasets now publicly available. Careful consideration of the confounders in these datasets is a critical step in the multi-omics pipeline. Enhanced understanding of the strengths and limitations of current and emerging multi-omic analytic approaches, and of the advantages of sparse machine learning pipelines to overcome current barriers in BPD research, are important contributions provided by the above pilot project.

In conclusion, cord blood multi-omics provides a snapshot at birth of the complex interplay of the perinatal metabolome, proteome and exposome that contribute to the pathogenesis of BPD, mediated by preterm birth. Integration of other emerging omics, such as genomics, transcriptomics, microbiomics in larger, multi-center studies with diverse biospecimen sources and serial sampling will further enhance our understanding of multifactorial BPD and its endotypes. Coupled with novel machine learning approaches, these future studies will refine our biomarker profiles for prediction and management of BPD, towards mitigation of the “perfect storm” which is critically needed for BPD prevention.

## Data Availability Statement

The datasets generated and analyzed for the current study are publicly available in https://doi.org/10.6084/m9.figshare.33147644.

## Acknowledgements

We thank the patients and families for their contributions and participation in this study. We thank additional members of the study team for their contributions and meticulous implementation of study protocols necessary for completing this project: Juanita Saqibuddin, RN, Kelly Stephens, RN, Rob Birkett, MSRC, Yeunook Bae, PhD.

## Author Contributions

K.M. and G.B. drafted the initial manuscripts, W.F., J.R. and I.S. contributed expertise in the manuscript drafts, preparation and interpretation of results. B.W., G.B., X.D., and J.H. designed, performed, drafted and prepared the results of machine learning and multi-omics analyses. K.M., J.N., J.Z., A.C. and W.F. and J.R. contributed to study design, data collection methods, cord blood assays and data annotation. K.M. conceived and designed the study, supervised the research, and obtained funding. All authors contributed to manuscript preparation and approved the final version.

## Consent Statement

Informed consent was obtained from all subjects involved in the parent study. The study was conducted in accordance with the Declaration of Helsinki, and the protocol was approved by the Institutional Review Board of Northwestern University (protocol number STU00201858).

