## Supplementary material for "Sparse Machine Learning Pipeline with Stabl Identifies Cord Blood Multi-Omic Signatures of Bronchopulmonary Dysplasia": Data Supplement File A

### Data Supplement File A: Selection of Features by LASSO and Stabl

### A1. Preterm vs. Full-term, Maximized Dataset

#### Metabolomics

| (# Stabl order) Selected By ALasso | Frequency of selection on CV | Model coefficient | p-value (Mann-Whitney) | Fold-Change |
|---|---|---|---|---|
| 1. plasma_2021909* | 0 | 0.48071 | 6.84E-22 | 1.79E+00 |
| 2. plasma_2012098* | 0 | 0.63523 | 9.17E-23 | 1.75E+00 |
| 3. plasma_2003510* | 0 | 0.56616 | 1.22E-23 | 1.84E+00 |
| 6. plasma_2010399* | 0 | 0.20423 | 6.89E-24 | 1.82E+00 |
| 7. plasma_2002997* | 0 | 0.08537 | 1.56E-13 | 1.14E+00 |
| 8. plasma_2033902* | 0 | 0.59376 | 4.74E-23 | 1.77E+00 |
| 11. plasma_2008333* | 0 | 0.61513 | 2.63E-24 | 1.87E+00 |
| 18. plasma_2021119* | 0 | 0.15836 | 6.47E-21 | 1.40E+00 |
| 19. plasma_2015417* | 0 | 0.09718 | 1.64E-21 | 1.74E+00 |

*Selected by ALasso and Stabl

#### Adductomics (Known)

| Selected by ALasso | Frequency of selection on CV | Model coefficient | (# order) Selected by Stabl | Frequency of selection on CV | Model coefficient | p-value (Mann-Whitney) | Fold-Change |
|---|---|---|---|---|---|---|---|
| Cys34 sulfinic acid plus methylation (not Cys34) | 1 | 1.38865 | 1. Cys34 sulfinic acid plus methylation (not Cys34) | 0.44 | 1.71776 | 7.96E-07 | 5.44E-01 |
| -Lys from C-terminus | 1 | -3.8538 | 2. -Lys from C-terminus | 0.08 | -8.06382 | 1.91E-07 | -1.29E-01 |
| S-Addition of hCys (NH2â†'OH) | 1 | 0.78803 | 3. S-Addition of hCys (NH2â†'OH) | 0.2 | 0.73438 | 2.72E-02 | 1.96E-01 |
| Methylation (not at Cys34) | 1 | -0.77285 | 4. Methylation (not at Cys34) | 0.04 | -1.13503 | 6.34E-02 | -1.99E-01 |
| Cys34 sulfonic acid (trioxidation) | 1 | 0.1577 | | | | 1.71E-04 | 5.88E-01 |
| S-Addition of crotonaldehyde | 1 | -0.00878 | | | | 2.95E-02 | -3.39E-01 |
| S-Addition of pyruvate or malonate semialdehyde | 1 | 0.28761 | | | | 5.94E-02 | 2.69E-01 |
| S-Addition of CysGly | 1 | 0.29279 | | | | 2.03E-01 | 1.37E-01 |
| dehydrated form of Cys34 sulfonic acid (trioxidation) | 1 | -0.01335 | | | | 7.53E-01 | -7.80E-02 |
| S-Methylthiolation_1 | 1 | -0.31468 | | | | 6.74E-01 | -9.43E-02 |
| S-Cys (possibly NH2 â†' OH, -H2O) | 1 | -0.37474 | | | | 1.39E-05 | -2.65E-02 |
| S-Methylethyl-sulfonylation | 1 | -0.21971 | | | | 2.21E-01 | -2.91E-02 |
| S-Addition of CysGly (-H2O) | 1 | -0.91612 | | | | 8.47E-01 | -9.50E-02 |
| K Adduct of S-CysGly | 1 | 0.15918 | | | | 2.49E-01 | 6.17E-02 |

### Adductomics (Unknown)

| Selected by ALasso | Frequency of selection on CV | Model coefficient | (# order) Selected by Stabl | Frequency of selection on CV | Model coefficient | p-value (Mann-Whitney) | Fold-Change |
|---|---|---|---|---|---|---|---|
| Unknown (153_05 Da) | 1 | -1.97767 | 1. Unknown (153_05 Da) | 1 | -3.21961 | 1.56E-06 | -4.75E-01 |
| Unknown (-10_07 Da) | 1 | 0.6296 | 2. Unknown (-10_07 Da) | 0.84 | 1.31334 | 3.95E-05 | 5.10E-01 |
| Unknown (138_06 Da) | 1 | 0.60795 | 3. Unknown (138_06 Da) | 0.2 | 0.65643 | 7.84E-03 | 2.95E-01 |
| Unknown (101_06 Da) | 1 | 0.64056 | | | | 2.60E-04 | 3.08E-01 |
| Unknown (109_03 Da) | 1 | -0.22208 | | | | 6.15E-01 | 3.13E-02 |
| Unknown (143 Da) | 1 | -1.50671 | | | | 6.35E-04 | -2.72E-01 |
| Unknown (180_02 Da) | 1 | 0.25314 | | | | 7.73E-02 | 9.21E-02 |
| Unknown (185_02 Da) | 1 | -0.35212 | | | | 2.49E-02 | -6.86E-02 |
| Unknown (212_32 Da) | 1 | -0.06117 | | | | 4.26E-02 | -8.00E-02 |
| Unknown (346_14 Da) | 1 | 0.49584 | | | | 2.16E-04 | 3.43E-01 |
| Unknown (351_07 Da) | 1 | -0.99427 | | | | 5.19E-02 | -7.92E-02 |
| Unknown (388_2 Da) | 1 | 0.68593 | | | | 4.33E-02 | 7.12E-02 |
| Unknown (461_2 Da) | 1 | 0.41313 | | | | 9.94E-01 | 1.35E-02 |
| Unknown (-12_96 Da), 2nd | 1 | -0.51709 | | | | 7.42E-03 | -1.84E-01 |
| Unknown (+340_08 Da), 2nd | 1 | -0.10846 | | | | 2.64E-03 | -9.17E-02 |
| Unknown (+545_22 Da), 2nd | 1 | -0.48149 | | | | 2.54E-03 | -2.52E-01 |

### A2. BPD on Preterm, Maximized Dataset

#### Metabolomics

| (# Stabl order) Selected By ALasso | Frequency of selection on CV | Model coefficient | p-value (Mann-Whitney) | Fold-Change |
|---|---|---|---|---|
| 1. plasma_2008918* | 0.16 | 0.09788 | 7.08E-04 | 7.31E-01 |
| 2. plasma_2032736* | 0.52 | 0.10733 | 3.85E-06 | 6.93E-01 |
| 3. plasma_2018738* | 0.48 | 0.1206 | 2.30E-06 | 6.66E-01 |
| 4. plasma_2031319* | 0.2 | 0.12988 | 1.29E-06 | 7.89E-01 |
| 5. plasma_1024124* | 0.32 | 0.0827 | 3.46E-04 | 5.29E-01 |
| 6. plasma_1021917* | 0.28 | 0.06324 | 4.02E-04 | 5.54E-01 |
| 7. plasma_1024253* | 0.28 | 0.1004 | 2.59E-05 | -6.92E-01 |
| 8. plasma_1023840* | 0.08 | 0.07008 | 1.52E-04 | -5.08E-01 |
| 9. plasma_2013663* | 0.04 | 0.07433 | 2.25E-03 | 2.69E-01 |
| 10. plasma_2000889* | 0.12 | 0.07959 | 3.33E-04 | 7.46E-01 |
| 11. plasma_1012076* | 0.2 | 0.07395 | 6.26E-04 | 5.62E-01 |

*Selected by ALasso and Stabl

#### Adductomics (Known and Unknown)

| (# Stabl order) Selected By ALasso | Omic | Frequency of selection on CV | p-value (Mann-Whitney) | Fold-Change |
|---|---|---|---|---|
| 1. S-Addition of hCys (-H2O) | Known | 0 | 3.71E-01 | -1.74E-02 |
| 2. Methylation (not at Cys34) | Known | 0 | 3.60E-01 | -1.49E-01 |
| 3. S-Methylethyl-sulfonylation | Known | 0 | 7.00E-01 | -3.63E-03 |
| 4. S-Phenylation | Known | 0 | 4.26E-01 | -1.65E-03 |
| 5. S-Cys (possibly NH2 â†' OH, -H2O) | Known | 0 | 9.49E-01 | 2.96E-03 |
| 6. Cys34 sulfinic acid plus methylation (not Cys34) | Known | 0 | 1.86E-01 | 2.30E-01 |
| 7. S-Addition of S2O3H | Known | 0 | 9.27E-01 | 1.07E-02 |
| 8. Cys34â†'Oxoalanine or formylglycine | Known | 0 | 5.17E-01 | -1.74E-05 |
| 9. S-hCys, plus methylation (not Cys34) | Known | 0 | 1.38E-01 | -1.16E-01 |
| 10. S-Addition of tiglic aldehyde | Known | 0 | 5.64E-01 | 1.08E-02 |
| 1. Unknown (126_08 Da) | Unknown | 0.16 | 4.24E-02 | 1.54E-01 |
| 2. Unknown (+476 Da) | Unknown | 0.08 | 1.59E-01 | 1.85E-01 |
| 3. Unknown (+509_21 Da) | Unknown | 0 | 2.18E-01 | -2.08E-02 |
| 4. Unknown (151_99 Da) | Unknown | 0 | 2.99E-01 | -2.02E-01 |
| 5. Unknown (111_03 Da) | Unknown | 0.12 | 9.48E-01 | -3.93E-03 |
| 6. Unknown (388_2 Da) | Unknown | 0 | 1.14E-01 | 8.05E-02 |
| 7. Unknown (+154_35 Da) | Unknown | 0 | 8.96E-01 | -1.29E-03 |
| 8. Unknown (62_01 Da) | Unknown | 0 | 6.09E-02 | -2.91E-02 |
| 9. Unknown (34_92 Da) | Unknown | 0 | 5.21E-01 | 3.34E-03 |
| 10. Unknown (+495_21 Da, M49,OS39) | Unknown | 0 | 7.03E-01 | -1.30E-02 |

### A3. BPD on Preterm, Reduced Dataset

#### Metabolomics

| Selected by ALasso | Frequency of selection on CV | Model coefficient | (# order) Selected by Stabl | Frequency of selection on CV | Model coefficient | p-value (Mann-Whitney) | Fold-Change |
|---|---|---|---|---|---|---|---|
| plasma_1005387 | 0.2 | 0.54507 | 1. plasma_1005387 | 0.4 | 3.51637 | 2.32E-05 | 1.31E+00 |
| plasma_2016415 | 0.2 | 0.13012 | 2. plasma_2016415 | 0.08 | 3.1858 | 8.91E-04 | 1.37E+00 |
| plasma_1023419 | 0.2 | -0.52088 | 3. plasma_1023419 | 0.04 | -3.53487 | 1.44E-03 | -9.88E-01 |
| plasma_2032915 | 0 | 0.34347 | 4. plasma_2032915 | 0.16 | 3.61639 | 1.65E-03 | 9.18E-01 |
| plasma_1026130 | 0.2 | 0.52518 | 5. plasma_1026130 | 0.12 | 4.67467 | 3.75E-04 | 1.00E+00 |
| plasma_1020047 | 0.2 | -0.22899 | | | | 2.86E-03 | -1.13E+00 |
| plasma_2022601 | 0.2 | -0.26331 | | | | 9.89E-04 | -9.99E-01 |
| plasma_2031044 | 0 | -0.36197 | | | | 6.95E-04 | -9.47E-01 |
| plasma_2038120 | 0 | -0.68199 | | | | 4.51E-04 | -1.02E+00 |

#### Adductomics (Known)

| Selected by ALasso | Frequency of selection on CV | Model coefficient | (# order) Selected by Stabl | Frequency of selection on CV | Model coefficient | p-value (Mann-Whitney) | Fold-Change |
|---|---|---|---|---|---|---|---|
| S-Phenylation | 1 | 3.56916 | 1. S-Phenylation | 0.96 | 3.27698 | 6.40E-03 | 4.13E-01 |
| Putative S-addition of acrolein | 0.76 | 0.56024 | 2. Putative S-addition of acrolein | 0.36 | 0.68372 | 7.32E-01 | 1.73E-03 |
| Cys34 sulfinic acid (dioxidation) | 0.96 | -2.527 | 3. Cys34 sulfinic acid (dioxidation) | 0.28 | -1.10727 | 7.18E-01 | 3.19E-02 |
| K Adduct of S-CysGly | 0.96 | 1.12816 | 4. K Adduct of S-CysGly | 0.64 | 1.30796 | 2.06E-02 | 4.11E-01 |
| S-Addition of SO2 | 0.8 | -0.93138 | 5. S-Addition of SO2 | 0.28 | -1.28916 | 7.77E-01 | 2.41E-01 |
| S-Methylthiolation | 0.52 | 0.17369 | 6. S-Methylthiolation | 0.48 | 0.26876 | 3.84E-01 | -6.00E-02 |
| | | | 7. dehydrated form of Cys34 sulfinic acid plus methylation (not Cys34) | 0.4 | -0.5459 | 3.05E-01 | -1.14E-04 |
| Methylation (not at Cys34) | 0.32 | -1.20118 | 8. Methylation (not at Cys34) | 0 | -0.13287 | 6.74E-01 | 3.15E-01 |
| Cys34â†’Gly | 0.44 | 0.70205 | | | | 1.62E-01 | 4.04E-01 |
| CH2 crosslink | 0.52 | 0.5001 | | | | 2.79E-02 | 7.80E-01 |
| Oxindole | 0.36 | 0.49355 | | | | 6.89E-01 | 1.83E-02 |
| Na adduct of S-Cys | 0.08 | 0.78905 | | | | 2.07E-01 | 1.57E-01 |
| S-(N-acetyl)Cys | 0.28 | -0.0595 | | | | 8.07E-01 | -8.05E-03 |
| S-CysGly, plus methylation (not Cys34) | 0.4 | -0.99969 | | | | 8.53E-01 | -4.77E-02 |
| dehydrated form of Cys34 sulfonic acid (trioxidation) | 0.4 | 0.53226 | | | | 3.24E-01 | 1.47E-01 |
| S-Addition of hCys (NH2â†’OH) | 0.28 | 1.0132 | | | | 4.76E-01 | -6.13E-02 |

### Adductomics (Unknown)

| (# order) Selected by Stabl | Frequency of selection on CV | Model coefficient | p-value (Mann-Whitney) | Fold-Change |
|---|---|---|---|---|
| 1. Unknown (+322_08 Da), 2nd | 0.16 | -2.68745 | 0.929918 | 0.007861 |
| 2. Unknown (101_06 Da) | 0.12 | 1.54756 | 0.252902 | 0.059346 |
| 3. Unknown (143 Da) | 0.16 | -0.23185 | 0.134883 | 0.399557 |
| 4. Unknown (360_19 Da) | 0.04 | -1.45073 | 0.261104 | -0.26872 |
| 5. Unknown (+509_21 Da) | 0.08 | 2.18176 | 0.776879 | -0.04903 |
| 6. Unknown (-12_96 Da) | 0.16 | -4.32993 | 0.120243 | -0.1215 |
| 7. Unknown (34_92 Da) | 0.12 | 30.6988 | 0.406186 | -0.00031 |
| 8. Unknown (+489 Da) | 0.16 | -1.35795 | 0.174364 | -0.28331 |
| 9. Unknown (62_01 Da) | 0.12 | -5.14817 | 0.100619 | -0.01686 |
| 10. Unknown (126_08 Da) | 0.32 | 3.25068 | 0.295742 | 0.082405 |
| 11. Unknown (156_96 Da) | 0.08 | -1.25239 | 0.041116 | 0.332328 |
| 12. Unknown (+495_21 Da, M49,OS39) | 0.08 | 2.39574 | 0.822168 | 0.020997 |
| 13. Unknown (262_1 Da) | 0.16 | -0.77795 | 0.286807 | -0.20728 |

### Proteomics

| Selected by ALasso | Frequency of selection on CV | Model coefficient | (# order) Selected by Stabl | Frequency of selection on CV | Model coefficient | p-value (Mann-Whitney) | Fold-Change |
|---|---|---|---|---|---|---|---|
| LACRT | 0.6 | 0.5682 | 1. LACRT | 0.44 | 2.15294 | 3.56E-02 | 2.83E-02 |
| SPDL1 | 0.44 | -0.49356 | 2. SPDL1 | 0.72 | -2.23759 | 1.50E-02 | -3.82E-01 |
| | | | 3. TNNT2 | 0.64 | -3.30623 | 1.67E-02 | -9.18E-01 |
| LRRC3B | 0.2 | 0.40741 | 4. LRRC3B | 0.12 | 4.56172 | 5.68E-03 | 6.30E-01 |
| TBCA | 0.44 | 0.72189 | 5. TBCA | 0.32 | 1.03266 | 9.24E-04 | 1.32E+00 |
| CDH12 | 0.48 | 0.59118 | 6. CDH12 | 0.32 | 2.04048 | 3.74E-02 | 2.77E-01 |
| RBL1 | 0.52 | -0.51548 | 7. RBL1 | 0.36 | -0.91659 | 3.70E-03 | -6.56E-01 |
| PLA2G2D | 0.24 | 0.41657 | 8. PLA2G2D | 0.2 | 2.81335 | 3.48E-02 | 2.28E-01 |
| EGFL6 | 0.56 | -0.76743 | 9. EGFL6 | 0.2 | -3.44623 | 1.11E-01 | -1.59E-01 |
| NPPB | 0.28 | -0.54547 | 10. NPPB | 0.4 | -3.35293 | 2.69E-01 | -1.63E-01 |
| | | | 11. IFNA10 | 0.48 | -1.57987 | 6.60E-01 | -4.53E-02 |
| | | | 12. DIPK1C | 0.12 | 2.2812 | 6.40E-03 | 1.06E+00 |
| | | | 13. DEFB135 | 0.28 | 3.75039 | 8.09E-03 | 1.07E+00 |
| LRPAP1 | 0.4 | -0.04904 | 14. LRPAP1 | 0.2 | -1.5896 | 1.58E-02 | -5.76E-01 |
| | | | 15. CA12 | 0.16 | 4.28492 | 1.35E-01 | 1.89E-01 |
| | | | 16. MCM6 | 0.08 | -1.6613 | 8.20E-02 | -2.79E-01 |
| | | | 17. TFPI | 0.24 | -1.79594 | 1.08E-02 | -6.97E-01 |
| | | | 18. TES | 0.12 | 0.53661 | 4.95E-02 | 5.00E-01 |
| UTS2R | 0.24 | -0.15095 | 19. UTS2R | 0.2 | -0.87758 | 6.48E-02 | -3.44E-01 |
| | | | 20. ATG4B | 0.24 | 1.08351 | 4.17E-01 | 5.35E-02 |
| | | | 21. ICAM4 | 0.28 | 0.23626 | 4.73E-02 | 5.12E-01 |
| | | | 22. RRM2B | 0.12 | 2.42963 | 3.74E-02 | 4.67E-01 |
| | | | 23. TLR10 | 0.12 | -0.04554 | 1.49E-03 | 7.46E-01 |
| | | | 24. IL4I1 | 0.04 | 1.09594 | 9.45E-01 | 2.05E-02 |
| | | | 25. HS6ST2 | 0.12 | 1.03056 | 3.74E-02 | 5.09E-01 |
| | | | 26. OAT | 0.12 | -2.27638 | 5.93E-02 | -4.83E-01 |
| | | | 27. ZNF334 | 0.16 | 1.84292 | 1.16E-01 | 2.05E-01 |
| | | | 28. NT5DC3 | 0.28 | -0.94102 | 5.04E-03 | -6.47E-01 |
| | | | 29. MGAT4C | 0.12 | 0.04072 | 9.86E-02 | 1.89E-01 |
| F13A1, F13B | 0.84 | 1.32902 | | | | 3.35E-04 | 1.35E+00 |
| CFD | 0.2 | 0.01851 | | | | 2.88E-03 | 8.85E-01 |
| CRISP2 | 0.36 | 0.66114 | | | | 2.65E-02 | 9.66E-01 |
| POLR2C | 0.12 | 0.31352 | | | | 1.94E-01 | 4.06E-03 |
| CBLN1 | 0.32 | -0.36408 | | | | 1.40E-01 | -6.00E-01 |
| ARF3 | 0.16 | 0.1406 | | | | 3.74E-01 | 9.14E-02 |
| DCUN1D2 | 0.36 | 0.02513 | | | | 1.77E-01 | 2.57E-01 |
| WWC1 | 0.24 | 0.25308 | | | | 3.33E-01 | 1.46E-01 |

### A4. BPD severity, Grade III vs. Rest, Maximized dataset

#### Metabolomics

| (# order) Selected By Stabl | Frequency of selection on CV | Model coefficient | p-value (Mann-Whitney) | Fold-Change |
|---|---|---|---|---|
| 1. plasma_1022095* | 0.2 | 0.05074 | 2.46E-01 | 5.09E-01 |
| 2. plasma_1010553 | 0.2 | 0.02108 | 3.03E-01 | 4.20E-02 |
| 3. plasma_1012499* | 0.4 | 0.03504 | 8.64E-04 | 8.28E-01 |
| 4. plasma_1005067 | 0 | 0.03907 | 9.47E-02 | -3.45E-01 |
| 5. plasma_2032502* | 0.2 | 0.03493 | 1.39E-03 | -6.26E-01 |
| 6. plasma_1003769 | 0 | 0.03715 | 2.69E-02 | 6.17E-01 |
| 7. plasma_2033888* | 0.4 | 0.03319 | 4.96E-01 | 1.44E-01 |
| 8. plasma_2021700* | 0.2 | 0.04272 | 6.99E-03 | 3.19E-01 |
| 9. plasma_1014836 | 0 | 0.04521 | 2.87E-03 | -6.79E-01 |
| 10. plasma_2016274 | 0.2 | 0.03059 | 6.76E-01 | 1.02E-01 |
| 11. plasma_2001288 | 0 | 0.01571 | 4.69E-01 | 3.87E-03 |
| 12. plasma_2001309 | 0 | 0.01925 | 4.74E-01 | -1.93E-03 |
| 13. plasma_1016900 | 0 | 0.02941 | 2.87E-03 | -4.39E-01 |
| 14. plasma_1011518 | 0 | 0.02047 | 2.75E-01 | -9.51E-02 |
| 15. plasma_2001352* | 0 | 0.04314 | 6.42E-03 | -7.27E-01 |
| 16. plasma_2020418 | 0.2 | 0.07063 | 2.92E-01 | -2.10E-02 |
| 17. plasma_2035249* | 0 | 0.02594 | 4.61E-01 | -1.28E-01 |
| 18. plasma_2008185 | 0 | 0.02022 | 3.09E-01 | -3.46E-01 |
| 19. plasma_1012982 | 0.2 | 0.01016 | 4.27E-01 | -4.16E-02 |
| 20. plasma_2036199* | 0 | 0.05844 | 4.14E-01 | 5.15E-01 |
| 21. plasma_2037388* | 0 | 0.02632 | 6.51E-03 | -8.09E-01 |
| 22. plasma_1016492 | 0.2 | 0.04684 | 2.75E-04 | 8.16E-01 |
| 23. plasma_2003517* | 0.2 | 0.02482 | 1.71E-01 | 1.00E-01 |
| 24. plasma_2038949* | 0 | 0.03019 | 1.62E-02 | 6.77E-01 |
| 25. plasma_2038644 | 0 | 0.02212 | 7.62E-02 | -3.41E-01 |
| 26. plasma_2027982 | 0 | 0.02296 | 3.92E-04 | 7.75E-01 |
| 27. plasma_2039130* | 0.2 | 0.03252 | 1.62E-02 | -6.59E-01 |
| 28. plasma_1010946 | 0 | 0.03493 | 2.31E-04 | -5.88E-01 |
| 29. plasma_2030953* | 0 | 0.01205 | 3.10E-01 | 1.13E-01 |
| 30. plasma_1005337* | 0 | 0.03156 | 7.94E-01 | 1.93E-01 |
| 31. plasma_2009324 | 0 | 0.03258 | 1.13E-04 | 9.04E-01 |

*Selected by Lasso and Stabl

#### Adductomics (Known)

| Selected by Lasso | Frequency of selection on CV | Model coefficient | (# order) Selected by Stabl | Frequency of selection on CV | Model coefficient | p-value (Mann-Whitney) | Fold-Change |
|---|---|---|---|---|---|---|---|
| S-Methylethyl-sulfonylation | 1 | -0.96999 | 1. S-Methylethyl-sulfonylation | 1 | 1 | 5.80E-01 | 6.81E-03 |
| Cys34â†'Gly | 1 | -0.47087 | | | | 1.59E-01 | 1.84E-01 |
| CH2 crosslink | 1 | 0.7655 | | | | 7.92E-01 | -3.31E-01 |
| Methylation (not at Cys34) | 1 | 0.49694 | | | | 2.25E-01 | -2.14E-01 |
| Cys34-Gln cross-link (monooxidation), Cys34 Sulfinamide | 1 | -0.10202 | | | | 3.03E-01 | -3.27E-01 |
| S-Sodiation | 1 | -1.22631 | | | | 2.77E-01 | -1.87E-01 |
| Ethylene oxide adduct | 1 | -0.40463 | | | | 9.90E-01 | 5.47E-02 |
| Sâ€"(O)â€"Oâ€"CH3 | 1 | 0.10912 | | | | 1.97E-01 | -1.12E-01 |
| Cys34 sulfonic acid (trioxidation) | 1 | -0.25651 | | | | 3.38E-01 | -4.77E-02 |
| Methylisocyanate adduct | 1 | 0.40476 | | | | 8.02E-01 | -2.46E-01 |
| S-Addition of SO2 | 1 | 0.76117 | | | | 3.47E-02 | -3.35E-01 |
| S-Addition of crotonaldehyde | 1 | -1.25457 | | | | 8.10E-01 | -1.65E-01 |
| S-Addition of pyruvate or malonate semialdehyde | 1 | -1.33787 | | | | 4.41E-01 | 1.47E-01 |
| S-Addition of mercaptoacetic acid | 1 | -0.12902 | | | | 7.81E-01 | 2.40E-01 |
| S-Addition of S2O3H | 1 | 0.29013 | | | | 4.16E-01 | 9.01E-03 |
| S-Addition of hCys (-H2O) | 1 | 2.25018 | | | | 2.19E-01 | -1.36E-02 |
| S-Cys | 1 | -2.88934 | | | | 9.55E-01 | -1.68E-02 |
| S-Addition of BDE | 1 | -0.01536 | | | | 1.65E-01 | -1.26E-01 |
| Oxindole | 1 | -0.17742 | | | | 7.07E-01 | 1.02E-01 |
| S-hCys, plus methylation (not Cys34) | 1 | 0.05786 | | | | 5.06E-01 | 1.30E-01 |
| S-(N-acetyl)Cys | 1 | 0.43149 | | | | 7.59E-01 | -1.77E-01 |
| S-Addition of CysGly | 1 | 1.23478 | | | | 1.33E-01 | -4.41E-01 |
| S-Addition of GluCys | 1 | 0.95019 | | | | 8.66E-01 | -1.80E-02 |
| S-Addition of GSH | 1 | -0.57397 | | | | 9.08E-01 | -2.68E-02 |
| -Lys from C-terminus | 1 | 1.68805 | | | | 6.01E-01 | -2.33E-02 |
| Cys34â†'Dehydroalanine | 1 | -0.08856 | | | | 1.91E-01 | -3.33E-03 |
| dehydrated form of Cys34 sulfonic acid (trioxidation) | 1 | -0.73056 | | | | 4.79E-01 | 2.22E-01 |
| S-Methylthiolation_1 | 1 | 1.40068 | | | | 2.51E-01 | -2.01E-01 |
| S-Methylthiolation_2 | 1 | 1.28384 | | | | 3.61E-01 | 3.72E-02 |
| Na adduct of S-CysGly | 1 | 0.02216 | | | | 2.37E-01 | -5.93E-02 |
| K adduct of T3 | 1 | 0.87344 | | | | 7.77E-02 | -2.84E-01 |

**Adductomics (Unknown)**

| (# order) Selected by Stabl | Frequency of selection on CV | Model coefficient | p-value (Mann-Whitney) | Fold-Change |
|---|---|---|---|---|
| 1. Unknown (126_08 Da) | 0.6 | 0.18166 | 6.92E-02 | 2.05E-02 |
| 2. Unknown (212_32 Da) | 0.8 | 0.16105 | 1.62E-01 | 1.66E-01 |
| 3. Unknown (+509_21 Da) | 1 | 0.25634 | 8.56E-01 | -1.10E-01 |
| 4. Unknown (138_06 Da) | 0.4 | 0.16644 | 7.33E-01 | 4.42E-01 |
| 5. Unknown (202_02 Da) | 0.4 | 0.23451 | 6.31E-01 | -1.19E-03 |

### A5. BPD severity, Grade IIIA vs Rest, Maximized dataset

#### Metabolomics

| (# order) Selected By Stabl | Frequency of selection on CV | Model coefficient | p-value (Mann-Whitney) | Fold-Change |
|---|---|---|---|---|
| 1. plasma_1022095 | 0.2 | 0.05074 | 2.30E-06 | -8.86E-01 |
| 2. plasma_1010553* | 0.2 | 0.02108 | 5.36E-01 | -4.20E-02 |
| 3. plasma_1012499 | 0.4 | 0.03504 | 2.40E-03 | -1.01E+00 |
| 4. plasma_1005067* | 0 | 0.03907 | 9.09E-01 | 2.13E-01 |
| 5. plasma_2032502 | 0.2 | 0.03493 | 2.39E-04 | 9.11E-01 |
| 6. plasma_1003769 | 0 | 0.03715 | 2.11E-01 | 3.16E-01 |
| 7. plasma_2033888 | 0.4 | 0.03319 | 1.62E-04 | -5.82E-01 |
| 8. plasma_2021700 | 0.2 | 0.04272 | 2.48E-04 | -9.09E-01 |
| 9. plasma_1014836 | 0 | 0.04521 | 2.67E-01 | -1.96E-01 |
| 10. plasma_2016274* | 0.2 | 0.03059 | 2.34E-01 | -1.31E-01 |
| 11. plasma_2001288* | 0 | 0.01571 | 4.67E-01 | -4.80E-03 |
| 12. plasma_2001309 | 0 | 0.01925 | 3.86E-01 | -2.70E-02 |
| 13. plasma_1016900 | 0 | 0.02941 | 8.38E-01 | 7.88E-02 |
| 14. plasma_1011518* | 0 | 0.02047 | 8.44E-01 | -1.06E-02 |
| 15. plasma_2001352 | 0 | 0.04314 | 2.98E-05 | 8.68E-01 |
| 16. plasma_2020418* | 0.2 | 0.07063 | 8.29E-02 | -2.60E-01 |
| 17. plasma_2035249 | 0 | 0.02594 | 2.35E-04 | 8.61E-01 |
| 18. plasma_2008185* | 0 | 0.02022 | 2.24E-01 | -2.81E-01 |
| 19. plasma_1012982* | 0.2 | 0.01016 | 7.68E-01 | 6.66E-02 |
| 20. plasma_2036199 | 0 | 0.05844 | 8.61E-04 | -9.23E-01 |
| 21. plasma_2037388 | 0 | 0.02632 | 3.77E-04 | 7.13E-01 |
| 22. plasma_1016492 | 0.2 | 0.04684 | 1.90E-04 | -6.72E-01 |
| 23. plasma_2003517 | 0.2 | 0.02482 | 1.94E-04 | -6.17E-01 |
| 24. plasma_2038949 | 0 | 0.03019 | 4.69E-04 | -8.47E-01 |
| 25. plasma_2038644 | 0 | 0.02212 | 2.83E-01 | 1.63E-01 |
| 26. plasma_2027982 | 0 | 0.02296 | 2.34E-02 | -6.55E-01 |
| 27. plasma_2039130 | 0.2 | 0.03252 | 1.71E-03 | 7.56E-01 |
| 28. plasma_1010946 | 0 | 0.03493 | 3.76E-01 | 9.39E-02 |
| 29. plasma_2030953 | 0 | 0.01205 | 2.08E-03 | -4.87E-01 |
| 30. plasma_1005337 | 0 | 0.03156 | 1.46E-03 | -5.14E-01 |
| 31. plasma_2009324 | 0 | 0.03258 | 8.90E-02 | -2.61E-01 |

*Selected by Lasso and Stabl

#### Adductomics (Known)

| Selected by Lasso | Frequency of selection on CV | Model coefficient | (# order) Selected by Stabl | Frequency of selection on CV | Model coefficient | p-value (Mann-Whitney) | Fold-Change |
|---|---|---|---|---|---|---|---|
| | | | 1. S-Methylethyl-sulfonylation | 1 | 1 | 1.26E-01 | 1.11E-02 |
| CH2 crosslink | 1 | -2.92908 | | | | 3.58E-01 | 3.35E-01 |
| Cys34 sulfinic acid (dioxidation) | 1 | 0.55255 | | | | 6.35E-01 | 1.07E-01 |
| Ethylene oxide adduct | 1 | -1.29616 | | | | 7.13E-01 | 1.19E-01 |
| Sâ€“(O)â€“Oâ€“CH3 | 1 | 3.19596 | | | | 8.89E-01 | 1.24E-02 |
| S-Methylthiolation | 1 | -0.0156 | | | | 5.18E-01 | 1.35E-01 |
| Cys34 sulfonic acid (trioxidation) | 1 | 1.1276 | | | | 5.65E-01 | -5.78E-03 |
| Na adduct of Cys34 sulfinic acid | 1 | -0.67743 | | | | 6.36E-01 | -8.09E-03 |
| S-Addition of SO2 | 1 | -1.22659 | | | | 8.38E-03 | 4.09E-01 |
| S-Addition of crotonaldehyde | 1 | 1.23964 | | | | 4.73E-01 | 5.06E-02 |
| S-Addition of tiglic aldehyde | 1 | -1.5513 | | | | 9.63E-01 | -1.30E-01 |
| S-Addition of Cys (-H2O) | 1 | 0.63738 | | | | 9.80E-01 | 5.33E-02 |
| S-Addition of hCys (-H2O) | 1 | -0.58647 | | | | 3.21E-02 | 3.46E-02 |
| S-CysGly, plus methylation (not Cys34) | 1 | -1.9395 | | | | 6.38E-02 | 1.45E-01 |
| S-Addition of GluCys | 1 | -0.29518 | | | | 4.09E-01 | 4.16E-02 |
| Cys34â†'Dehydroalanine | 1 | -0.20968 | | | | 4.84E-01 | 4.71E-03 |
| Cys34â†'Oxoalanine or formylglycine | 1 | 0.40382 | | | | 5.62E-01 | 6.98E-04 |
| dehydrated form of Cys34 sulfonic acid (trioxidation) | 1 | 0.14527 | | | | 8.43E-01 | -2.00E-01 |
| S-Methylthiolation_1 | 1 | -0.57771 | | | | 1.89E-01 | 6.14E-01 |
| S-Methylthiolation_2 | 1 | -2.33041 | | | | 6.13E-02 | 1.76E-01 |

### Adductomics (Unknown)

| (# order) Selected by Stabl | Frequency of selection on CV | Model coefficient | p-value (Mann-Whitney) | Fold-Change |
|---|---|---|---|---|
| 1. Unknown (126_08 Da) | 0.6 | 0.18166 | 8.99E-02 | -2.88E-02 |
| 2. Unknown (212_32 Da) | 0.8 | 0.16105 | 5.93E-02 | -1.52E-01 |
| 3. Unknown (+509_21 Da) | 1 | 0.25634 | 1.36E-01 | -1.67E-01 |
| 4. Unknown (138_06 Da) | 0.4 | 0.16644 | 6.92E-01 | -1.71E-01 |
| 5. Unknown (202_02 Da) | 0.4 | 0.23451 | 1.13E-01 | 6.53E-03 |

### A6. BPD Severity Grade III, Reduced Dataset

#### Metabolomics

| (# order) Selected by Stabl | Frequency of selection on CV | Model coefficient | p-value (Mann-Whitney) | Fold-Change |
|---|---|---|---|---|
| 1. plasma_1026126* | 0 | 12.41697 | 6.05E-03 | -1.07E+00 |
| 2. plasma_2007224* | 0.13333 | 2.95791 | 6.04E-02 | -6.69E-01 |
| 3. plasma_2034751 | 0.06667 | 4.04517 | 2.28E-04 | 1.29E+00 |
| 4. plasma_2030843* | 0.06667 | 5.0318 | 1.11E-02 | -9.49E-01 |
| 5. plasma_2037388* | 0 | 4.01776 | 3.74E-03 | -9.30E-01 |
| 6. plasma_2004555 | 0 | -3.49582 | 1.63E-04 | 1.46E+00 |
| 7. plasma_1002233* | 0 | 7.56412 | 8.19E-04 | -1.27E+00 |

#### Adductomics (Known)

| Selected by ALasso | Frequency of selection on CV | Model coefficient | (# order) Selected by Stabl | Frequency of selection on CV | Model coefficient | p-value (Mann-Whitney) | Fold-Change |
|---|---|---|---|---|---|---|---|
| S-Methylethyl-sulfonylation | 1 | -0.78079 | 1. S-Methylethyl-sulfonylation | 0.93333 | -1.29915 | 3.54E-01 | 1.58E-03 |
| Cys34-Dehydroalanine | 1 | 0.82281 | 2. Cys34-Dehydroalanine | 0.86667 | 1.03493 | 1.24E-01 | -1.41E-01 |
| CH2 crosslink | 1 | -0.46793 | | | | 1.05E-01 | 2.81E-01 |
| Cys34 sulfonic acid (trioxidation) | 1 | -0.76797 | | | | 9.94E-01 | -8.71E-03 |
| S-Addition of tiglic aldehyde | 1 | -2.22322 | | | | 1.49E-01 | 4.77E-01 |
| S-Addition of pyruvate or malonate semialdehyde | 1 | -0.13845 | | | | 2.31E-01 | 4.31E-01 |
| S-addition of benzaldehyde or quinone methide | 1 | -0.6086 | | | | 3.61E-01 | 3.22E-01 |
| S-Addition of S2O3H | 1 | 0.31917 | | | | 9.34E-01 | 8.72E-02 |
| S-Addition of BDE | 1 | 0.45766 | | | | 8.89E-01 | 3.02E-02 |
| S-Addition of CysGly | 1 | 0.49399 | | | | 8.52E-02 | -6.78E-01 |
| S-CysGly, plus methylation (not Cys34) | 1 | 0.1046 | | | | 3.11E-01 | -3.22E-02 |
| S-Addition of GluCys | 1 | 1.00864 | | | | 6.92E-01 | 2.51E-01 |
| -Lys from C-terminus | 1 | 0.06942 | | | | 7.30E-02 | -1.67E-01 |
| dehydrated form of Cys34 sulfinic acid plus methylation (not Cys34) | 1 | -0.8924 | | | | 7.38E-01 | -4.13E-05 |
| S-Methylthiolation_1 | 1 | 1.8619 | | | | 5.34E-01 | -4.08E-02 |
| S-Addition of CysGly (-H2O) | 1 | -0.00323 | | | | 8.42E-01 | 1.41E-01 |

#### Adductomics (Unknown)

#### None

### Proteomics:

| Selected by ALasso | Frequency of selection on CV | Model coefficient | (# order) Selected by Stabl | Frequency of selection on CV | Model coefficient | p-value (Mann-Whitney) | Fold-Change |
|---|---|---|---|---|---|---|---|
| CD33 | 0.8 | 0.67743 | 1. CD33 | 0.33333 | 1.78319 | 1.12E-02 | -1.42E+00 |
| SCN3B | 0.6 | 0.14867 | 2. SCN3B | 0.26667 | 0.97824 | 2.69E-03 | -1.07E+00 |
| | | | 3. IL18R1 | 0.2 | 0.3351 | 9.12E-04 | -1.34E+00 |
| NCR1 | 0.33333 | 0.06485 | | | | 1.29E-01 | -2.78E-01 |
| CGA, CGB3, CGB7 | 0.33333 | -0.19942 | | | | 3.72E-01 | 2.52E-01 |
| CD5 | 0.33333 | 0.01451 | | | | 8.91E-02 | -8.80E-01 |
| PRSS2 | 0.4 | 0.42957 | | | | 7.53E-01 | -3.91E-01 |
| UBC | 0.2 | -0.03601 | | | | 1.12E-01 | 6.28E-01 |
| PRKAR1A | 0.53333 | 0.18965 | | | | 1.07E-02 | -6.83E-01 |
| PSG2 | 0.26667 | -0.14251 | | | | 1.15E-01 | 1.39E-01 |
| LCP1 | 0.26667 | 0.10161 | | | | 2.96E-01 | -6.42E-02 |
| SELL | 0.2 | 0.06827 | | | | 8.86E-01 | -1.35E-01 |
| IDO1 | 0.33333 | 0.0657 | | | | 2.36E-01 | 9.32E-02 |
| ITIH1 | 0.33333 | -0.01024 | | | | 3.95E-01 | 4.50E-01 |
| TNFRSF1B | 0.6 | 0.23051 | | | | 3.38E-02 | -4.18E-01 |
| BGN | 0.53333 | -0.04522 | | | | 7.59E-01 | 1.47E-01 |
| PGM1 | 0.66667 | 0.4912 | | | | 7.49E-02 | -5.38E-01 |
| HK2 | 0.2 | 0.00073 | | | | 2.97E-01 | 1.76E-01 |
| BASP1 | 0.4 | 0.23627 | | | | 4.16E-02 | -8.27E-01 |
| PSG9 | 0.66667 | -0.10654 | | | | 6.04E-03 | 1.06E+00 |
| RAPGEF1 | 0.2 | 0.02055 | | | | 3.41E-01 | -5.34E-01 |
| PSG3 | 0.46667 | -0.26549 | | | | 3.89E-02 | 4.83E-01 |
| LRRN1 | 0.46667 | 0.21431 | | | | 1.04E-01 | -1.21E+00 |
| LYSMD3 | 0.33333 | -0.18856 | | | | 4.28E-01 | 1.03E-01 |
| IRX2-DT | 0.33333 | 0.36792 | | | | 1.78E-02 | -1.01E+00 |
| LILRB2 | 0.66667 | 0.6361 | | | | 2.71E-02 | -8.18E-01 |
| LRTM2 | 0.4 | 0.18536 | | | | 6.35E-02 | -8.05E-01 |
| LILRB1 | 0.8 | 0.54247 | | | | 1.20E-02 | -6.57E-01 |
| LRRC20 | 0.33333 | -0.03064 | | | | 1.55E-01 | 6.59E-01 |
| BRK1 | 0.33333 | -0.10697 | | | | 1.59E-01 | 4.48E-01 |
| TADA1 | 0.2 | -0.07807 | | | | 7.31E-01 | -2.94E-01 |
| S100A13 | 0.33333 | -0.18881 | | | | 8.07E-01 | 1.12E-01 |
| HEPH | 0.33333 | -0.15077 | | | | 5.34E-02 | 6.74E-02 |
| REPIN1 | 0.53333 | 0.05068 | | | | 1.16E-01 | -1.16E+00 |
| SEMA4A | 0.2 | 0.02975 | | | | 2.80E-01 | -3.10E-01 |
| SAP30L | 0.46667 | 0.2713 | | | | 1.11E-01 | -6.12E-01 |
| IL1RL2 | 0.53333 | -0.00825 | | | | 2.46E-02 | 5.99E-01 |
| RBM22 | 0.46667 | -0.01608 | | | | 8.22E-01 | 7.34E-01 |
| NXT1 | 0.8 | 0.85495 | | | | 1.94E-01 | -2.24E-01 |
| CFAP45 | 0.4 | 0.23458 | | | | 2.81E-01 | -3.56E-01 |
| VSIG4 | 0.46667 | 0.07183 | | | | 5.10E-02 | -8.57E-01 |
| CSDC2 | 0.53333 | 0.13937 | | | | 6.50E-02 | -1.04E+00 |
| PCDHGA1 | 0.33333 | -0.02139 | | | | 1.93E-01 | 6.90E-01 |
