## Supplementary material for "Sparse Machine Learning Pipeline with Stabl Identifies Cord Blood Multi-Omic Signatures of Bronchopulmonary Dysplasia": Data Supplement File B

**Data Supplement: Confounder Analysis Tables**

**Table A1:** Confounder analysis for preterm birth in relation to different clinical features. a. Difference of the summary statistics between the primary model, containing only cross-validated prediction, and the extended models, containing the prediction with the confounders. b. Coefficients significance of confounders in the model with confounders.

| Feature | Coef | Std err | Z | P>\|z\| |
|---|---|---|---|---|
| a. Difference (D) between models with and without confounders | | | | |
| **Pred** | Δ = -2.96 | Δ = 0.05 | Δ = -17.2 | Δ = 0.0 |
| b. Confounders coefficient | | | | |
| **Pred** | -1,050 | 0,039 | -26,939 | 0,000 |
| **apgar_5 minute** | -0,399 | 0,035 | -11,260 | 0,000 |
| **Birthweight** | -1,213 | 0,036 | -33,278 | 0,000 |
| **GA** | -1,221 | 0,033 | -37,364 | 0,000 |
| **PTL** | 0,691 | 0,043 | 15,890 | 0,000 |
| **apgar_1 minute** | -0,430 | 0,059 | -7,327 | 0,000 |
| **Preeclampsia** | 0,351 | 0,059 | 5,944 | 0,000 |
| **mode_of_delivery** | -0,387 | 0,076 | -5,115 | 0,000 |
| **antenatal_steroids** | 0,201 | 0,041 | 4,848 | 0,000 |
| **race_White** | -0,263 | 0,057 | -4,625 | 0,000 |
| **maternal_age** | -0,238 | 0,055 | -4,299 | 0,000 |
| **bw_percentile** | 0,267 | 0,063 | 4,269 | 0,000 |
| **multiple_gestation_status_Twin** | 0,154 | 0,064 | 2,401 | 0,016 |
| **clinical_chorio** | 0,090 | 0,040 | 2,245 | 0,025 |
| **race_Black or African American** | 0,110 | 0,050 | 2,212 | 0,027 |
| **multiple_gestation_status_Triplet** | 0,055 | 0,029 | 1,879 | 0,060 |
| **race_Unknown** | 0,025 | 0,015 | 1,683 | 0,092 |
| **rupture of membrane_type** | 0,093 | 0,075 | 1,235 | 0,217 |
| **race_Other** | 0,063 | 0,059 | 1,065 | 0,287 |
| **race_Asian** | 0,045 | 0,065 | 0,693 | 0,488 |
| **Ethnicity** | 0,054 | 0,082 | 0,655 | 0,512 |
| **race_Declined** | 0,028 | 0,043 | 0,649 | 0,516 |
| **race_Hispanic** | -0,002 | 0,003 | -0,618 | 0,536 |
| **gender_coded_0male_1female** | -0,000 | 0,064 | -0,004 | 0,997 |

**Table A2.** Confounder analysis for BPD in relation to different clinical features with the maximized dataset. a. Difference of the summary statistics between the primary model, containing only the cross-validation prediction, and the extended models, containing the prediction with the confounders. b. Coefficients significance of confounders in the model with confounders.

| Feature | Coef | Std err | Z | P>\|z\| |
|---|---|---|---|---|
| a. Difference (D) between models with and without confounders | | | | |
| **Pred** | Δ = -0.19 | Δ = 0.09 | Δ = -1.73 | Δ = 0.003 |
| b. Confounders coefficient | | | | |
| **Pred** | -0,311 | 0,265 | -1,172 | 0,241 |
| **birthweight** | -1,051 | 0,334 | -3,146 | 0,002 |
| **race_Unknown** | 0,469 | 0,176 | 2,671 | 0,008 |
| **multiple_gestation_status_Triplet** | -0,424 | 0,170 | -2,493 | 0,013 |
| **rupture of membrane_type** | 0,627 | 0,255 | 2,456 | 0,014 |
| **antenatal_steroids** | 0,452 | 0,266 | 1,697 | 0,090 |
| **race_Hispanic** | 0,174 | 0,109 | 1,603 | 0,109 |
| **ga** | -0,497 | 0,326 | -1,527 | 0,127 |
| **race_Black or African American** | -0,435 | 0,345 | -1,260 | 0,208 |
| **preeclampsia** | 0,336 | 0,290 | 1,161 | 0,246 |
| **ethnicity** | 0,437 | 0,436 | 1,003 | 0,316 |
| **gender_coded_0male_1female** | -0,218 | 0,278 | -0,781 | 0,435 |
| **apgar_1 minute** | -0,212 | 0,314 | -0,674 | 0,500 |
| **race_Declined** | -0,221 | 0,342 | -0,645 | 0,519 |
| **apgar_5 minute** | -0,221 | 0,344 | -0,641 | 0,521 |
| **mode_of_delivery** | -0,148 | 0,280 | -0,527 | 0,598 |
| **maternal_age** | 0,107 | 0,268 | 0,401 | 0,688 |
| **multiple_gestation_status_Twin** | 0,070 | 0,242 | 0,291 | 0,771 |
| **PTL** | -0,078 | 0,305 | -0,255 | 0,799 |
| **race_White** | 0,096 | 0,380 | 0,252 | 0,801 |
| **clinical_chorio** | -0,075 | 0,311 | -0,242 | 0,809 |
| **race_Other** | 0,077 | 0,402 | 0,190 | 0,849 |
| **bw_percentile** | 0,031 | 0,302 | 0,102 | 0,919 |
| **race_Asian** | 0,018 | 0,289 | 0,061 | 0,952 |

**Table A3.** Confounder analysis for BPD in relation to different clinical features with the reduced dataset. a. Difference of the summary statistics between the primary model, containing only the cross validation prediction, and the extended models, containing the prediction with the confounders. b. Coefficients significance of confounders in the model with confounders.

| Feature | Coef | Std err | Z | P>\|z\| |
|---|---|---|---|---|
| a. Difference (D) between models with and without confounders | | | | |
| **Pred** | Δ = -0.19 | Δ = 0.09 | Δ = -1.73 | Δ = 0.003 |
| b. Confounders coefficient | | | | |
| **Pred** | -0,495 | 0,409 | -1,212 | 0,226 |
| **ga** | -0,845 | 0,244 | -3,468 | 0,001 |
| **rupture of membrane_type** | 0,776 | 0,318 | 2,438 | 0,015 |
| **apgar_1 minute** | -0,705 | 0,328 | -2,147 | 0,032 |
| **multiple_gestation_status_Triplet** | -0,337 | 0,201 | -1,675 | 0,094 |
| **race_Unknown** | 0,177 | 0,120 | 1,476 | 0,140 |
| **race_Black or African American** | -0,495 | 0,353 | -1,403 | 0,160 |
| **race_Other** | -0,473 | 0,374 | -1,264 | 0,206 |
| **clinical_chorio** | -0,351 | 0,304 | -1,156 | 0,248 |
| **multiple_gestation_status_Twin** | -0,427 | 0,393 | -1,088 | 0,276 |
| **bw_percentile** | -0,310 | 0,295 | -1,048 | 0,295 |
| **maternal_age** | 0,340 | 0,336 | 1,013 | 0,311 |
| **race_Declined** | -0,235 | 0,262 | -0,896 | 0,370 |
| **gender_coded_0male_1female** | -0,282 | 0,354 | -0,797 | 0,426 |
| **ethnicity** | -0,176 | 0,451 | -0,392 | 0,695 |
| **preeclampsia** | 0,102 | 0,315 | 0,323 | 0,746 |
| **apgar_5 minute** | -0,124 | 0,413 | -0,300 | 0,764 |
| **PTL** | -0,030 | 0,335 | -0,090 | 0,928 |
| **mode_of_delivery** | -0,029 | 0,353 | -0,081 | 0,936 |
| **race_White** | 0,013 | 0,400 | 0,033 | 0,974 |
| **antenatal_steroids** | 1,178 | 0,267 | 4,417 | 0,000 |
| **birthweight** | -0,955 | 0,226 | -4,216 | 0,000 |

**Table A4.** Confounder analysis for BPD severity (III vs Rest) in relation to different clinical features with the maximized dataset. a. Difference of the summary statistics between the primary model, containing only the cross-validation prediction, and the extended models, containing the prediction with the confounders. b. Coefficients significance of confounders in the model with confounders.

| Feature | Coef | Std err | Z | P>\|z\| |
|---|---|---|---|---|
| a. Difference (D) between models with and without confounders | | | | |
| **Pred** | Δ = -0.19 | Δ = 0.09 | Δ = -1.73 | Δ = 0.003 |
| b. Confounders coefficient | | | | |
| **Pred** | 0,583 | 0,370 | 1,576 | 0,115 |
| **race_Declined** | -0,976 | 0,278 | -3,511 | 0,000 |
| **PTL** | 1,078 | 0,452 | 2,385 | 0,017 |
| **birthweight** | -0,594 | 0,322 | -1,843 | 0,065 |
| **race_Hispanic** | -0,258 | 0,157 | -1,642 | 0,101 |
| **clinical_chorio** | 0,568 | 0,367 | 1,551 | 0,121 |
| **race_Other** | -0,507 | 0,379 | -1,338 | 0,181 |
| **bw_percentile** | -0,431 | 0,330 | -1,306 | 0,192 |
| **rupture of membrane_type** | -0,450 | 0,397 | -1,133 | 0,257 |
| **preeclampsia** | -0,468 | 0,504 | -0,929 | 0,353 |
| **race_Unknown** | -0,253 | 0,299 | -0,848 | 0,396 |
| **apgar_1 minute** | 0,311 | 0,367 | 0,847 | 0,397 |
| **mode_of_delivery** | -0,274 | 0,371 | -0,739 | 0,460 |
| **maternal_age** | -0,270 | 0,391 | -0,690 | 0,490 |
| **multiple_gestation_status** | -0,178 | 0,369 | -0,482 | 0,630 |
| **ga** | 0,153 | 0,337 | 0,453 | 0,651 |
| **race_Black or African American** | -0,194 | 0,433 | -0,448 | 0,654 |
| **antenatal_steroids** | 0,085 | 0,343 | 0,249 | 0,804 |
| **apgar_5 minute** | -0,081 | 0,363 | -0,224 | 0,823 |
| **race_White** | -0,063 | 0,384 | -0,164 | 0,870 |
| **ethnicity** | 0,051 | 0,435 | 0,118 | 0,906 |
| **gender_coded_0male_1female** | 0,027 | 0,381 | 0,071 | 0,944 |

**Table A5.** Confounder analysis for BPD severity (IIIA vs Rest) in relation to different clinical features with the maximized dataset. a. Difference of the summary statistics between the primary model, containing only the cross-validation prediction, and the extended models, containing the prediction with the confounders. b. Coefficients significance of confounders in the model with confounders.

| Feature | Coef | Std err | Z | P>\|z\| |
|---|---|---|---|---|
| a. Difference (D) between models with and without confounders | | | | |
| **Pred** | Δ = -0.19 | Δ = 0.09 | Δ = -1.73 | Δ = 0.003 |
| b. Confounders coefficient | | | | |
| **Pred** | 0,932 | 0,368 | 2,530 | 0,011 |
| **maternal_age** | 1,102 | 0,401 | 2,746 | 0,006 |
| **mode_of_delivery** | 0,747 | 0,273 | 2,734 | 0,006 |
| **apgar_1 minute** | 0,693 | 0,268 | 2,587 | 0,010 |
| **race_Other** | -0,532 | 0,214 | -2,483 | 0,013 |
| **race_Hispanic** | 0,813 | 0,402 | 2,022 | 0,043 |
| **multiple_gestation_status** | -0,419 | 0,216 | -1,942 | 0,052 |
| **rupture of membrane_type** | 0,318 | 0,171 | 1,854 | 0,064 |
| **ethnicity** | -0,731 | 0,433 | -1,687 | 0,092 |
| **race_Unknown** | -0,101 | 0,065 | -1,568 | 0,117 |
| **race_White** | 0,379 | 0,247 | 1,536 | 0,124 |
| **PTL** | 0,293 | 0,191 | 1,531 | 0,126 |
| **race_Declined** | 0,644 | 0,427 | 1,509 | 0,131 |
| **preeclampsia** | 0,322 | 0,215 | 1,497 | 0,135 |
| **apgar_5 minute** | 0,354 | 0,305 | 1,163 | 0,245 |
| **clinical_chorio** | 0,320 | 0,341 | 0,939 | 0,348 |
| **gender_coded_0male_1female** | -0,200 | 0,273 | -0,733 | 0,464 |
| **race_Black or African American** | 0,231 | 0,369 | 0,627 | 0,531 |
| **ga** | -0,077 | 0,279 | -0,275 | 0,783 |
| **birthweight** | -0,067 | 0,260 | -0,258 | 0,796 |
| **antenatal_steroids** | -0,058 | 0,419 | -0,138 | 0,890 |
| **bw_percentile** | -0,026 | 0,292 | -0,088 | 0,930 |

**Table A6.** Confounder analysis for BPD severity (III vs Rest) in relation to different clinical features with the reduced dataset. a. Difference of the summary statistics between the primary model, containing only the cross-validation prediction, and the extended models, containing the prediction with the confounders. b. Coefficients significance of confounders in the model with confounders.

| Feature | Coef | Std err | Z | P>\|z\| |
|---|---|---|---|---|
| a. Difference (D) between models with and without confounders | | | | |
| **Pred** | Δ = -0.19 | Δ = 0.09 | Δ = -1.73 | Δ = 0.003 |
| b. Confounders coefficient | | | | |
| **Pred** | 1,253 | 0,267 | 4,699 | 0,000 |
| **PTL** | 0,841 | 0,207 | 4,051 | 0,000 |
| **preeclampsia** | -0,460 | 0,137 | -3,350 | 0,001 |
| **maternal_age** | -0,684 | 0,276 | -2,481 | 0,013 |
| **ethnicity** | 0,583 | 0,239 | 2,438 | 0,015 |
| **apgar_1 minute** | 0,801 | 0,331 | 2,418 | 0,016 |
| **mode_of_delivery** | -0,611 | 0,324 | -1,889 | 0,059 |
| **race_Other** | -0,232 | 0,135 | -1,719 | 0,086 |
| **bw_percentile** | -0,413 | 0,246 | -1,682 | 0,093 |
| **race_Unknown** | 0,332 | 0,221 | 1,503 | 0,133 |
| **clinical_chorio** | 0,466 | 0,311 | 1,496 | 0,135 |
| **apgar_5 minute** | -0,482 | 0,330 | -1,457 | 0,145 |
| **race_Declined** | -0,209 | 0,149 | -1,404 | 0,160 |
| **birthweight** | -0,261 | 0,234 | -1,119 | 0,263 |
| **race_Black or African American** | 0,229 | 0,316 | 0,724 | 0,469 |
| **race_White** | -0,174 | 0,297 | -0,588 | 0,557 |
| **multiple_gestation_status** | -0,161 | 0,307 | -0,526 | 0,599 |
| **gender_coded_0male_1female** | -0,172 | 0,333 | -0,517 | 0,605 |
| **rupture of membrane_type** | -0,070 | 0,329 | -0,212 | 0,832 |
| **antenatal_steroids** | -0,069 | 0,329 | -0,210 | 0,833 |
| **ga** | 0,032 | 0,259 | 0,123 | 0,902 |
